# Hv1 proton channel is essential for splenic antibacterial defense and immune homeostasis during the chronic phase of traumatic brain injury in male mice

**DOI:** 10.64898/2026.07.31.742029

**Authors:** Zhuofan Lei, Romeesa Khan, Yun Li, Kavitha Brunner, Cole R Sebok, Patrick J Devlin, Junyun He, Rodney M. Ritzel, Junfang Wu

**Affiliations:** Department of Anesthesiology & Center for Shock, Trauma and Anesthesiology Research (STAR), University of Maryland School of Medicine, Baltimore, MD 21201, USA; Department of Neurology, University of Texas Health Science Center at Houston, McGovern Medical School, Houston, TX 77030, USA; MD Anderson Cancer Center UTHealth Graduate School of Biomedical Sciences, Houston, TX 77030, USA; Center for Neuroimmunology and Glial Biology, Institute of Molecular Medicine, University of Texas Health Science Center at Houston, Houston, TX 77030, USA

**Keywords:** Traumatic brain injury, Hv1 proton channel, long-term survival, peripheral immune dysfunction, splenic antibacterial defense, neuroinflammation, transcriptomic profiling

## Abstract

**Background:** Traumatic brain injury (TBI) is increasingly recognized as a chronic condition with lasting systemic consequences. Beyond persistent neuroinflammation, long-term TBI disrupts peripheral immune homeostasis, increasing susceptibility to infection and organ dysfunction, particularly in older patients. The voltage-gated proton channel Hv1, expressed in microglia and peripheral immune cells, regulates oxidative injury via modulation of NADPH oxidase activity. Yet few animal studies extend long enough to recapitulate the lifelong trajectory of human TBI, leaving the long-term effects of Hv1 deficiency on systemic immune homeostasis unresolved.

**Methods:** Young adult (3-month-old) male wild-type (WT) and Hv1 knockout (Hv1KO) mice were subjected to a moderate controlled cortical impact (CCI), and survival was monitored for up to 18 months post-injury, with endpoint analyses performed at 21 months of age. After neurological behavioral assessments, spleen, lung, liver, gut, ipsilateral cortex, and blood samples were collected for flow cytometry, qPCR, NanoString nCounter Panels, and *in vivo* plasma transfer studies.

**Results:** Hv1 deficiency resulted in significantly increased mortality following TBI, starting at 14 months post-injury, compared with WT/TBI mice. No significant difference in survival was observed between the two sham groups. At 18 months post-injury, Hv1KO mice exhibited significant weight loss and splenomegaly. qPCR revealed an approximately 30-fold increase of pan-bacterial 16S rRNA levels in the spleens of Hv1KO/TBI mice, but not in the lungs or liver. Furthermore, chronic TBI in the Hv1KO mice led to a compromised intestinal tight junction and mucus barrier integrity, accompanied by aberrant activation of the cyclic GMP-AMP synthase-stimulator of interferon genes pathway in the spleen. Transcriptomic profiling of the spleen, liver, and lung revealed distinct post-injury immune signatures in Hv1KO mice. In contrast, surviving Hv1KO/TBI mice showed modest behavioral resilience and a partially neuroprotective cortical transcriptomic profile. Lastly, systemic transfer of plasma from WT/TBI or Hv1KO donors into naïve young adult mice altered immune responses in the spleen, lung, and brain.

**Conclusions:** Hv1 plays a critical role in maintaining peripheral immune integrity and antibacterial defense throughout the chronic course of TBI. Despite conferring modest neuroprotection through attenuation of microglial-mediated oxidative stress, Hv1 deficiency exacerbated systemic phagocyte dysfunction and significantly reduced long-term survival.

## Introduction

Traumatic brain injury (TBI) is a leading cause of long-term disability worldwide and a major risk factor for chronic neurodegeneration, cognitive decline, and premature mortality [1, 2]. Beyond the initial mechanical insult, TBI triggers a prolonged secondary injury cascade that extends far beyond the central nervous system (CNS), leading to persistent systemic inflammation, immune suppression, gut barrier dysfunction, and altered peripheral organ function that can persist for months to years [2–4]. These peripheral disturbances are especially relevant in older adults, who now constitute the fastest-growing segment of the TBI population [5, 6]. Because aging independently erodes immune competence, tissue repair, and barrier integrity [7–9], brain injury in this group is superimposed on a pre-existing loss of systemic resilience, leaving survivors more prone to poor recovery, maladaptive immune responses, and chronic complications.

The voltage-gated proton channel Hv1, encoded by *Hvcn1*, has been identified as a key mediator of pathological microglial activation and a potential therapeutic target [10–12]. Hv1 provides the charge compensation required to sustain NADPH oxidase 2 (NOX2)-mediated reactive oxygen species (ROS) production during the phagocyte respiratory burst [13]. Within the CNS, Hv1 is expressed predominantly by microglia [10]. In young adult mice, genetic ablation of Hv1 confers neuroprotection across multiple CNS injury models, including ischemic stroke [10, 14], spinal cord injury [15–17], and demyelinating disease [18]. In TBI, we previously demonstrated that microglial Hv1 sustains pathological acidosis and oxidative burst activity, and that Hv1-deficient young adult mice exhibit improved cognitive and motor functional outcomes for up to one year post-injury [11, 19], establishing Hv1 as a therapeutic target. However, Hv1 is broadly expressed across peripheral phagocyte compartments and immune lineages, including neutrophils, monocytes/macrophages, dendritic cells, eosinophils, basophils, and B lymphocytes [13, 20–23]. Given its role in regulating phagocyte oxidative burst, host defense, and inflammatory responses, age-related changes in Hv1 signaling could affect systemic immune homeostasis after TBI. Notably, both mRNA and protein expression of Hv1 in the CNS increase with aging and after SCI [24, 25], suggesting that Hv1-dependent mechanisms may become progressively more important in aged TBI survivors. Despite this, prior studies of Hv1 in TBI have predominantly focused on acute and subacute post-injury time windows in relatively young mice. The impact of Hv1 on peripheral immune homeostasis remains unexplored in the context of chronic TBI.

In this study, we investigated the long-term consequences of constitutive Hv1 deficiency for up to 18 months following contusion-induced brain injury, one of the longest follow-up periods reported for this gene and TBI model. Using multidisciplinary approaches, we found that Hv1 deficiency unexpectedly impaired systemic immune homeostasis, leading to reduced survival, peripheral immune dysfunction, splenomegaly, bacterial accumulation, gut barrier disruption, and dysregulated cyclic GMP-AMP synthase-stimulator of interferon genes (cGAS-STING) signaling. We also identified circulating plasma factors as drivers of persistent microgliosis and systemic immune responses. These findings highlight Hv1 as a critical regulator at the interface of neuroinflammation and systemic immunity, underscoring the need to carefully balance the therapeutic potential of Hv1 inhibition against its essential role in host defense.

## Materials and Methods

### Animals and controlled cortical impact model of TBI

Young adult (3-4-month-old) male wild-type (WT) and Hv1 knockout (Hv1KO) mice on a C57BL/6 background were used in all experiments. Mice were housed under a 12-hour light/dark cycle with ad libitum access to food and water. Moderate TBI was induced using a controlled cortical impact (CCI) device (Precision Systems and Instrumentation, PSI), as previously described [11]. Briefly, mice were anesthetized and placed in a stereotaxic frame, followed by the craniotomy performed over the left somatosensory cortex. The impactor was positioned above the exposed cortical surface and delivered a single impact with a deformation depth of 1.2 mm and a dwell time of 100 ms at a velocity of 4 m/s. Sham-operated animals underwent anesthesia, scalp incision, without craniotomy or impact. Following incision closure with sutures, mice were allowed to recover in a warmed cage until fully ambulatory and then returned to their home cages. Postoperative monitoring was performed daily for up to 18 months. All procedures were approved by the Institutional Animal Care and Use Committees (IACUC) of the University of Maryland School of Medicine and University of Texas Health Science Center at Houston.

### Behavioral testing

A battery of cognitive, social, and affective tests was conducted to investigate the neurological behavioral phenotypes of WT and Hv1KO mice before the endpoint after TBI.

### Open field (OF)

Spontaneous locomotor activity and general exploratory behavior were measured in an open-field arena [26, 27]. Mice were placed in the center of the arena and allowed to explore freely for 5 minutes, which were video recorded and analyzed by AnyMaze system. Locomotor activity was quantified by measuring the total distance traveled during the testing session. Anxiety-like behavior was evaluated by analyzing the proportion of time spent in the center versus the periphery of the arena, with increased time spent in the peripheral zone interpreted as increased anxiety-like behavior.

### Y-maze

Working memory and spatial exploration were assessed via a standard Y-maze as previously described [26, 27]. Each mouse was placed at the end of one arm and allowed to explore freely for 6 minutes, starting from the animal’s 1^st^ full entry into the central platform. The locomotion activities were recorded, and spontaneous alternation was calculated as the ratio of consecutive entries into all three arms to the total triads. This assay provided measures of both working memory (alternation) and general activity (total entries).

### Novel object recognition (NOR)

Recognition memory was evaluated using a three-phase NOR paradigm [26, 27]. Mice were placed in the apparatus for 5-minute habituation on the first day. In next acquisition phase, mice were exposed to two identical objects symmetrically placed in the arena and allowed to explore until a 30-second total exploration time or 15-minute session duration was reached; object bias at baseline was calculated from this phase. During the following choice phase, one random familiar object was replaced with a novel object, and exploration time directed toward each object was recorded under the identical time window. Novelty preference was calculated as the percentage of time spent exploring the novel object relative to total object exploration.

### Three-chamber social recognition (SR)

Social behavior was investigated in a self-made three-chamber apparatus adapted from standard social recognition protocols as previously described [27]. The test consisted of three sequential phases: habituation, social preference, and social novelty preference. During habituation, mice were allowed to freely explore all three empty chambers and two empty wire cups for 5 minutes. During the social preference phase, a novel conspecific (“stranger 1”) was placed under one random cup while the opposite cup remained empty; time spent exploring each cup was recorded. During the social novelty phase, a second novel mouse (“stranger 2”) was placed under another empty cup with the first stranger remained; social novelty preference was expressed as the percentage of time spent exploring the cup containing the novel mouse. This assay quantified sociability and social memory.

### Novelty-suppressed feeding (NSF)

Affective-like behavior was evaluated using the NSF test based on established paradigm [27]. After 24-hour food deprivation, mice were placed in a novel arena where a petri dish with food pellets was positioned in the center. Latency to begin feeding in the novel arena was recorded as an index of anxiety-like behavior, and latency to feed when subsequently returned to the home cage was recorded as a control for appetite and motivation.

### RNA isolation and NanoString assay

At 18 months post-injury, mice were euthanized, blood samples were collected using cardiac puncture, and the animals were perfused with 50 mL ice-cold saline. Total RNA was isolated from dissected spleen, lung, liver, gut, and ipsilateral cortical tissues using the RNeasy Mini Kit (Qiagen, Cat# 74106). RNA samples were subsequently submitted to the Institute for Genome Sciences at the University of Maryland School of Medicine for NanoString gene expression analysis [28]. Transcriptomic profiling was performed on spleen, lung, and liver samples using the NanoString Myeloid Innate Immunity Panel, while cortical samples were analyzed using the NanoString Neuropathology Panel. Briefly, equal amounts of RNA were hybridized with barcoded probe sets, processed on the nCounter system, and normalized using internal controls and housekeeping genes according to standard NanoString workflows. The transcriptomic data was further processed in NanoString nSolver software (Advance analysis, ver. 2.0.134) to produce gene abundance (normalized log2 values), and differential expressed genes (DEGs) between any two groups. Additionally, NanoString gene-expression data from spleen, lung, and liver were subjected to cell-type deconvolution to infer immune-cell enrichment scores for neutrophils, macrophages, B cells, cytotoxic cells, dendritic cells, and other populations, using panel-embedded signatures and manufacturer-recommended pipelines.

Further in-depth analyses were performed by self-programmed R (ver. 4.6.0) code in RStudio (Ver. 2026.04.0 Build 526). The normalized log2 gene abundance data was used for cross-group expression visualization (heatmaps and boxplots), Partial Least Squares Discriminant Analysis (PLS-DA), single-sample Gene Set Enrichment Analysis (ssGSEA) by NanoString panel, and factor-decomposition analyses. The DEGs data derived from between-group comparison or factor-decomposition analyses was used for volcano graphs, Venn diagrams, pathway enrichment analysis, panel-based GSEA, and Protein-protein Interaction Network (PIN). The ssGSEA scores and cell-type scores were analyzed statistically through two-way ANOVA and post hoc testing.

### Real Time quantitative PCR (RT-qPCR)

After equal amounts of the total RNA were converted to cDNA using a Verso cDNA kit (ThermoFisher, Cat# AB1453B), RT-qPCR reactions were run through thermal cycling consisted of an initial UNG incubation at 50°C for 2 min and AmpliTaq activation at 95°C for 10 min, followed by 40 cycles of denaturation at 95°C for 15 s and annealing/extension at 60°C for 1 min using TaqMan assays on a QuantStudio^TM^ 5 Real-Time PCR System (Applied Biosystems, Carlsbad, CA). Target mRNAs were measured using TaqMan gene expression assays (Supplemental table 1). Wells returning undetermined Ct values were assigned 0 to the final relative fold change. Relative mRNA expression was calculated by normalization to GAPDH with the WT Sham group as control using the 2^-ΔΔCt^ method [29].

### Flow cytometry

Flow cytometry sample preparation and cell staining were performed as reported previously [19, 30]. Briefly, mice were transcardially perfused with 20 mL of cold saline prior to removal of tissue. The ipsilateral brain hemisphere was collected and filtered through a 70-μm filter into a 50 mL conical tube. Cells were resuspended in complete Roswell Park Memorial Institute (RPMI) 1640 (Invitrogen, Cat# 22,400,105) medium and digested in collagenase/dispase (1 mg/mL, Roche, Cat# 50-100-3281), papain (5 U/mL suspension, Worthington Biochemical, Cat# LS003126), 0.5 mM EDTA (Research Products International, Cat# E14000), and DNAase I (10 mg/mL, Roche, Cat# 10104159001) for 35 min in a shaking incubator at 200 rpm and 37 °C. The cell suspension was washed with 5 mL of RPMI, centrifuged at 500 g for 5 min at 4 °C, then resuspended in a final volume of supplemented RPMI at 5 mL/hemisphere and kept on ice until staining.

For spleen processing, after filtration through a 70-μm filter into a 50 ml conical tube, cells were washed and treated with 3 ml of red blood cell lysis buffer (Thermo Scientific, Cat# 00-4333-57) for 10 min on ice, then splenocytes were washed and resuspended in 1 mL of supplemented RPMI and kept on ice until staining. Lung tissue (right superior and middle lobes) was collected into 5 mL of RPMI, mechanically chopped to a fine consistency, then digested in collagenase/dispase (0.5 mg/mL, Roche) and DNAase I (5 mg/mL, Roche) for 25 min at 200 rpm and 37 °C. Cells were then filtered through a 70-μm filter into 50 mL conical, washed, resuspended in 1 mL of supplemented RPMI, and kept on ice until staining.

All cells were distributed into FACS tubes for staining and washed with 1 mL of FACS buffer prior to staining. The cells were incubated with Fc block (Anti-CD16/32, BioLegend, Cat# 101302) for 10 min on ice prior to staining for the following surface antigens: CD45-violetFluor-450 (30-F11, Cytek Bio, Cat# 5-0451-U025), CD11b-APC/Fire™ 750 (M1/70, BioLegend, Cat# 101262), Ly6C-AF700 (HK1.4, BioLegend, Cat# 128024), Ly6G-Bv650 (1A8, BioLegend, Cat# 127641), MHCII-PerCP/Cy5.5 (M5/114.15.2, BioLegend, Cat# 107262). The fixable viability dye Zombie Aqua or Zombie UV (BioLegend, Cat# 423102 or 423108) was used for live/dead discrimination. Cells were then washed in FACS buffer, fixed in 2.0% paraformaldehyde for 10 min, and washed once more prior to adding 300 μL FACS buffer. Following 2 hours of brefeldin A incubation in supplemented RPMI, intracellular cytokine/protein staining was performed as described previously [31] using a fixation/permeabilization kit (BD Biosciences, Cat# 554714) and the following antibodies: CD68-PECy7 (FA-11, BioLegend, Cat# 137016), Syk-PE (F5F, BioLegend, Cat# 646004), NOX2-AF647 (Bioss Antibodies, bs-3889R), and TNF-Bv510 (MP6-XT22, BioLegend, Cat# 506339) per the manufacturer’s instructions. IgG bead uptake was performed using the Phagocytosis Assay Kit IgG-FITC (Cayman Chemical Company, Cat# 500290) as per the manufacturer’s instructions. To measure antigen processing and ovalbumin hydrolysis, the fluorescently conjugated DQ-Ovalbumin (Thermo Scientific, Cat# D12053) was used as per the manufacturer’s instructions. For ROS measurement, cells were loaded with the fluorescent probe dihydrorhodamine-123 (DHR123; ThermoFisher, Cat# D23806).

Cell counts were measured as absolute counts based on volumetric sampling or counting bead estimation (BioLegend, Cat# 424902). Leukocytes were first gated using a splenocyte reference (SSC-A vs FSC-A). Singlets were gated (FSC-H vs FSC-W), and live cells were gated based on Zombie Aqua exclusion (SSC-A vs Zombie Aqua-Bv510 or Zombie-UV). Resident microglia were identified as CD45^int^CD11b^+^Ly6C^−^, whereas peripheral leukocytes were distinguished as CD45^hi^CD11b^+^ myeloid cells and CD45^hi^CD11b^−^ lymphocytes. Myeloid cell populations were further divided into Ly6C^+^Ly6G^+^ neutrophils and Ly6C^hi^Ly6G^-^ monocytes. Cell type-matched fluorescence minus one (FMO) controls were used to determine the positivity of each antibody.

### Processing of donor plasma and *in vivo* plasma transfer injections

Donor mouse blood was collected using cardiac puncture. Immediately, cells were removed from the blood by centrifugation for 15 min at 12,000 rpm using a refrigerated (4 °C) centrifuge. The supernatant (plasma) was collected into new microcentrifuge tubes and pooled by group. The plasma was then dialyzed using the Slide-A-Lyzer dialysis cassettes with 3.5 kDa molecular weight cut-off (Thermo Fisher Scientific, Cat# 66330), for removal of EDTA (residual anticoagulant). The plasma was then aliquoted and stored at −80 °C until required for injections. For the plasma transfer experiments, young male WT (C57BL/6J) recipient mice were administered 200 µl of donor plasma via alternating retroorbital injections, from either aged WT/sham, aged WT/TBI, aged Hv1KO/sham, or aged Hv1KO/TBI donors. Plasma injections were administered once per day for 4 consecutive days. Injections were performed on deeply anesthetized mice and mice were monitored for post-injection recovery for 4 h after each injection.

### Statistical analysis

Survival curves were generated by the Kaplan-Meier method and compared between groups using log-rank (Mantel-Cox) tests, with additional Gehan-Breslow-Wilcoxon test for trend or weighted comparisons. Molecular, cellular, and behavioral data were analyzed by two-way ANOVA with genotype and injury as fixed factors; three-way ANOVA was used for selected behavioral experiments with additional within-subject factors (e.g., object or cup). When significant main or interaction effects were detected, Tukey’s or Sidak’s post hoc tests were applied as indicated in the figure legends. Statistical analyses were performed using GraphPad Prism (GraphPad Software, Inc., La Jolla, CA, ver. 8.4.2), with data presented as mean ± SEM. Outliers were determined using Grubb’s test. *p* ≤ 0.05 was considered statistically significant.

## Results

### Chronic TBI in Hv1-deficient mice reduces long-term survival and promotes spleen-restricted bacterial accumulation

To examine the long-term consequences of Hv1 deficiency after moderate TBI, young adult male WT and Hv1KO mice were subjected to CCI or sham surgery and monitored for 18 months. Hv1KO/TBI mice exhibited pronounced late-onset mortality beginning at 14 months post-injury (6 of 17 dead by the endpoint), whereas WT/TBI mice showed only a single death and both sham cohorts maintained 100% survival (**Fig. 1A**). Among survivors, body weight was lower with TBI in WT mice and with Hv1 deficiency in sham mice, without further exacerbation by the gene-injury interaction (**Fig. 1B**). Spleen weight was elevated in Hv1KO mice independently of injury, indicating genotype-driven splenomegaly (**Fig. 1C**). As the spleen is a major reservoir of NOX2-dependent phagocytes [32], we measured splenic bacterial burden by RT-qPCR for pan-bacterial 16S rRNA. Hv1KO/TBI spleens showed an approximately 30-fold elevation in 16S rRNA relative to WT/Sham (**Fig. 1D**), with no significant differences among other groups. Neither lung nor liver showed altered 16S signal across groups (**Fig. 1E-F**), indicating that the bacterial accumulation is spleen-restricted. Species-specific qPCR for Escherichia coli (E. coli) and Staphylococcus aureus (S. aureus) in all three organs was below detection threshold (**Supplemental Fig. 1**), excluding these classical post-traumatic pathogens [33, 34]. Together, these results reveal an unexpected long-term vulnerability in Hv1 deficiency mice after TBI, marked by delayed mortality, splenomegaly, and spleen restricted bacterial accumulation.

**Fig. 1.**
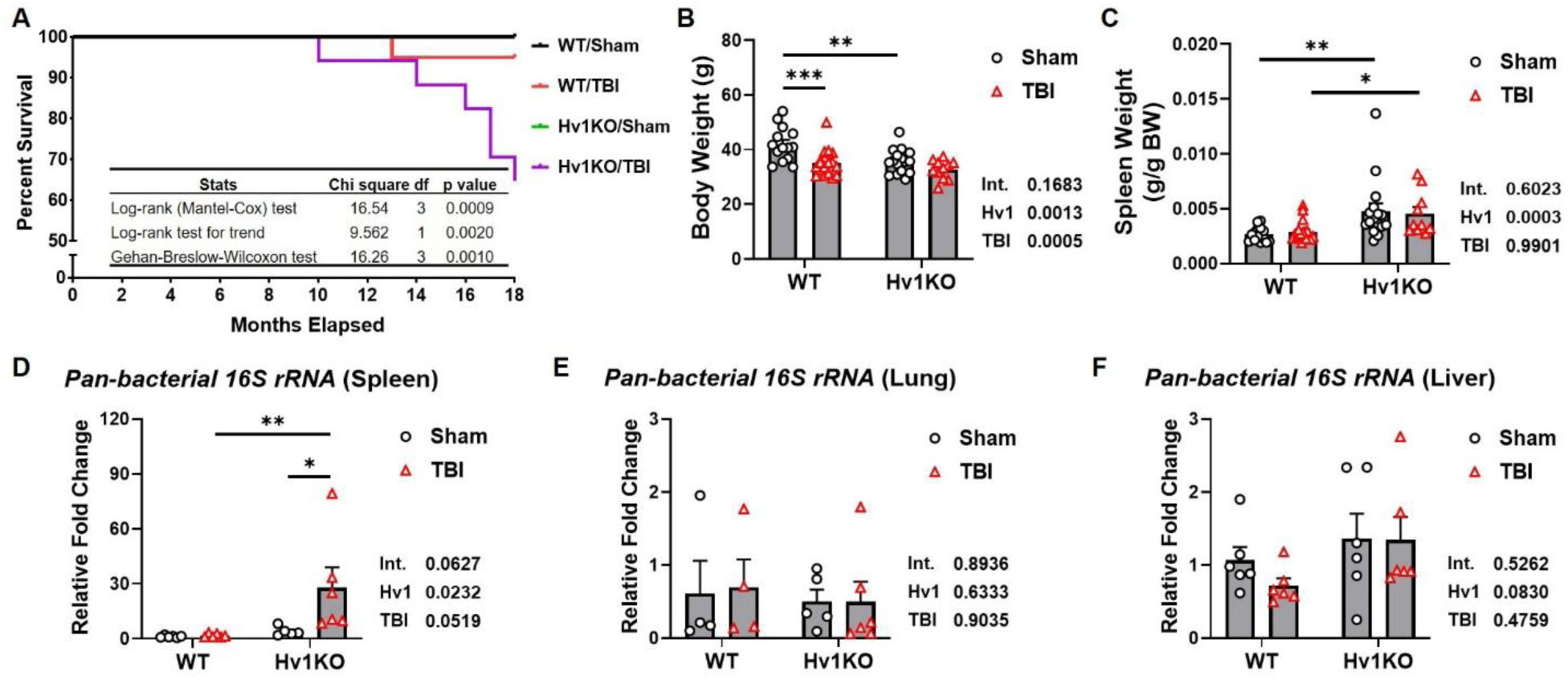
Hv1 deficiency is associated with reduced long-term survival and increased pan-bacterial 16S rRNA in the spleen at 18 months post-TBI. (A) Kaplan-Meier analysis showing reduced survival in Hv1KO/TBI mice relative to other groups. (B) Body weight was lower with TBI (in WT) and with Hv1 deficiency (in sham). (C) Hv1KO mice showed elevated spleen weight (splenomegaly) independently of injury. (D) Splenic pan-bacterial 16S rRNA was elevated in Hv1KO/TBI relative to WT/Sham and Hv1KO/Sham. (E-F) 16S signal did not differ across groups in lung (E) or liver (F). Sample sizes (A-C): WT/Sham n=16; Hv1KO/Sham n=16; WT/TBI n=18; Hv1KO/TBI n=11. Sample sizes (D-F): n=4-6 mice/group. Statistics: two-way ANOVA with Šídák-corrected simple-effects comparisons (B-F); interaction (Int.), Hv1, and TBI main-effect p-values shown beside each panel. *p≤0.05, **p≤0.01, ***p≤0.001, ****p≤0.0001.

### Hv1 deficiency drives constitutive remodeling of the splenic immune transcriptome that decompensates under chronic TBI

To investigate the molecular mechanisms underlying splenomegaly and splenic bacterial accumulation in Hv1KO mice after chronic TBI, we profiled spleen tissue using NanoString Myeloid Innate Immunity panel. Unsupervised hierarchical clustering demonstrated distinct gene signatures of each group, and PLS-DA (44.9% variance explained) segregated the spleen samples into four non-overlapping clusters, indicating that both Hv1 deficiency and TBI cause robust transcriptional impacts in the spleen (**Fig. 2A-B**).

**Fig. 2.**
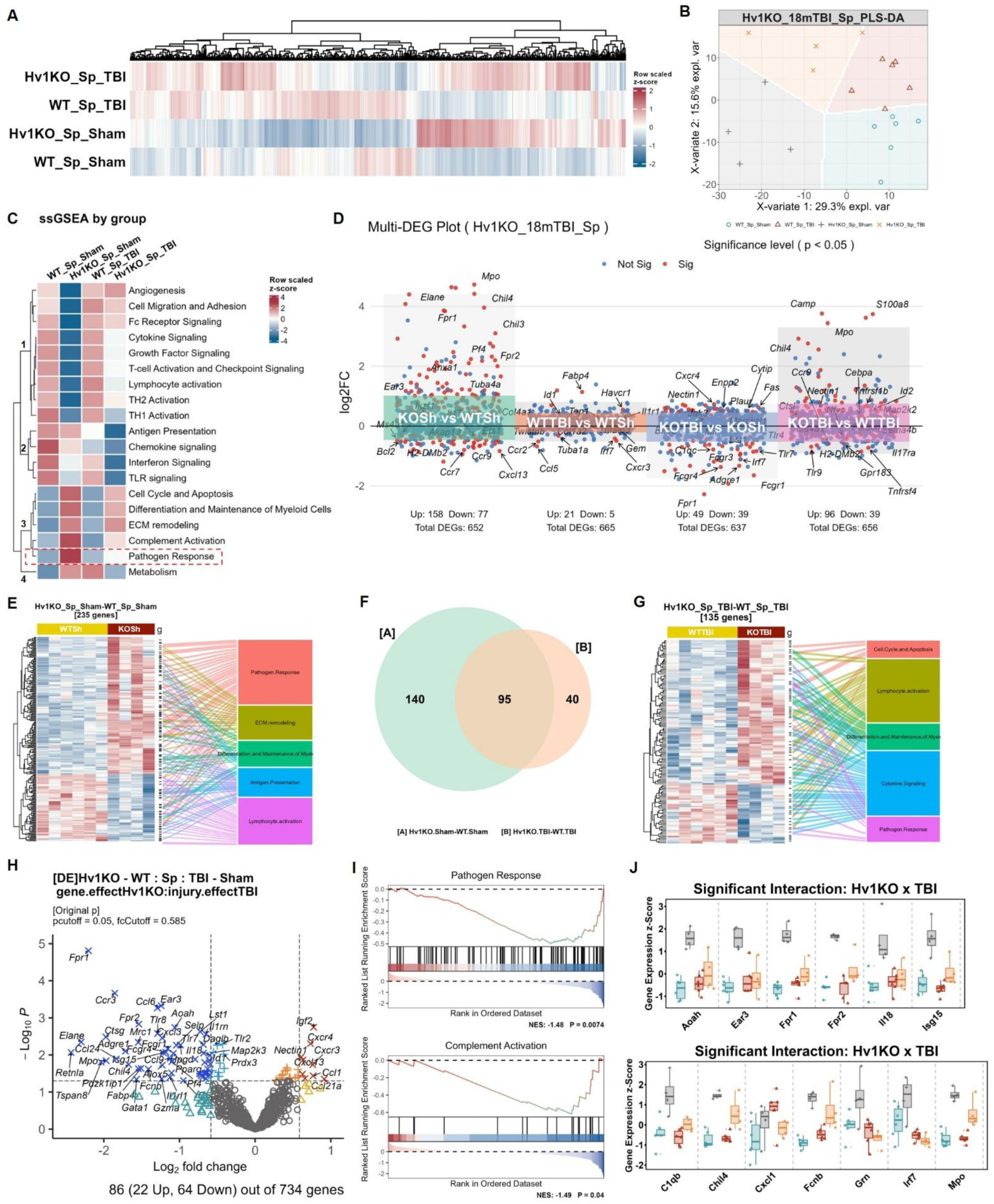
Hv1 deficiency in the spleen alters constitutive transcriptome and dysregulates immune responses to chronic TBI. **(A)** Myeloid Innate Immunity panel reveals distinct gene expression heatmaps from the spleen tissue of the four groups. **(B)** PLS-DA shows clear separation of transcriptomic signatures among groups. **(C)** The heatmap from ssGSEA of Myeloid Innate Immunity panel demonstrates robust changes induced by Hv1 deficiency with distinct post-injury responses. **(D)** DEGs between groups indicate up- and down-regulated transcriptional responses to Hv1 KO or TBI. **(E)** The expression profile of DEGs with Hv1KO/Sham versus WT/Sham is displayed in z-score heatmap, which mostly clusters into top 5 enriched pathways in the panel. **(F)** The Venn diagram shows DEG numbers induced by Hv1 KO in Sham groups and TBI groups respectively. **(G)** The z-score heatmap of gene expression with top 5 enriched pathways is plotted based on DEGs from Hv1KO/TBI versus WT/TBI. **(H)** The volcano graph shows DEGs from the interaction of Hv1 KO and TBI in splenic transcriptomic data. (**I**) Panel-based GSEA analysis demonstrated that the interaction of Hv1 KO and TBI induced down-regulated pathways of “Pathogen Response” and “Complement Activation”. **(J)** The expression profiles of genes with significant interaction effect from the two pathways are displayed as z-scores across groups. Sample sizes: WT/Sham, n=6; Hv1KO/Sham, n=4; WT/TBI, n=6; Hv1KO/TBI, n=4.

Using the functional pathways defined by NanoString Myeloid Innate Immunity panel, we performed ssGSEA on the splenic transcriptomic data and decomposed genotype and injury effects by two-way ANOVA. Unsupervised clustering of the scaled ssGSEA scores (**Fig. 2C**) resolved four modules, coordinately defining splenic immune state across groups, with ANOVA terms (**Supplemental Table 2**) provided as statistical support. Module 1: Nine pathways, comparable between two WT groups, were uniformly depressed in baseline Hv1KO spleen and partially recovered post-injury, including Angiogenesis, Cell Migration and Adhesion, Fc Receptor Signaling, Cytokine and Growth Factor Signaling, T-cell Activation and Checkpoint Signaling, Lymphocyte activation, Th1/2 Activation. The baseline changes indicated a weakened immune-supportive environment due to Hv1 deficiency, while the partial rebound after TBI, compared with the WT counterpart, implied a compensatory but incomplete re-engagement of these programs. Module 2: Four splenic pathways were specifically suppressed in Hv1KO/TBI group, including Antigen Presentation, Chemokine, Interferon, and TLR signaling. This module captured a more specific injury-triggered collapse, suggesting loss of coordinated innate sensing and antigen-handling capacity under chronic injury. Module 3: Five pathways, unchanged in WT, were robustly elevated in baseline Hv1KO spleen and degraded post-injury, including Cell Cycle and Apoptosis, Differentiation and Maintenance of Myeloid Cells, ECM remodeling, Complement Activation, Pathogen Response. This module revealed a chronically constitutive activation of myeloid cells in Hv1-deficient spleen that decompensated functionally post-injury. Here, Pathogen Response was the only pathway showing significant gene *x* injury interaction, the most elevated in Hv1KO/Sham and collapsed after chronic TBI. Module 4: One single pathway, Metabolism, was boosted either by TBI in WT and Hv1 deficiency in Sham animals, but fall back due to the two factors convergence, with no genotype, injury, or interaction term reaching significance.

Differential expression analysis quantified these shifts (**Fig. 2D**). Genotype effects dominated: 235 DEGs distinguished Hv1KO/Sham from WT/Sham and 135 distinguished Hv1KO/TBI from WT/TBI, whereas injury alone yielded only 26 DEGs in WT and 88 in Hv1KO. The 235 DEGs derived from baseline genotype effect clustered into Pathogen Response, ECM Remodeling, and Differentiation and Maintenance of Myeloid Cells (predominantly upregulated) and Antigen Presentation, Lymphocyte Activation (predominantly downregulated) (**Fig. 2E**). They shared 95 genes with the 135 DEGs between two injury groups (**Fig. 2F**). The latter clustered into a partially distinct top functional gene set (Cell Cycle and Apoptosis, Lymphocyte Activation, Differentiation/Maintenance of Myeloid Cells, Cytokine Signaling, Pathogen Response; **Fig. 2G**), indicating a TBI-driven shift in transcriptional response of Hv1 deficiency.

To extract genes driven by gene *x* injury interaction, we performed two-factor differential expression analysis, identifying 86 interaction-driven DEGs (22 up, 64 down; **Fig. 2H**). Prominent downregulation centered on bacterial sensing and killing-formyl peptide receptors *Fpr1*/*Fpr2*, neutrophil granule effectors *Elane*, *Mpo*, *Ctsg*, pattern recognition receptors *Tlr8*/*Tlr9*, and myeloid chemokines *Ccl6*, *Ccr3*. Panel-based GSEA on the interaction term confirmed suppression of Pathogen Response and Complement Activation (**Fig. 2I**). Cross-group expression profiles of genes with significant interaction from the two pathways demonstrated that the baseline elevation in Hv1KO/Sham was not sustained under chronic injury (**Fig. 2J**). PIN analysis identified ITGAM as the central hub linking C3AR1, SERPINB2/B6A, and inflammatory mediators (IL1B, IL6, TNF, CXCL1, CXCL10, NOS2, CHIL3, MPO) (**Supplemental Fig. 2A-B**), placing integrin- and complement-associated signaling at the molecular core of the decompensated splenic phenotype. Additionally, DEGs derived from genotype effect or injury effect were analyzed accordingly (**Supplemental Fig. 2C-F**), demonstrating dominant genotype effect at gene and pathway levels.

In summary, Hv1 loss remodels the splenic transcriptome bidirectionally, suppressing adaptive and immune-supportive programs while constitutively activating myeloid and pathogen-defense pathways with core genes disrupted. It fails to sustain antimicrobial output under chronic injury and links the transcriptomic phenotype to the splenomegaly and splenic bacterial accumulation.

### cGAS-STING-IFN signaling and pro-inflammatory responses are dysregulated in Hv1 KO spleen after chronic TBI

To test whether DNA-sensing pathways were engaged in response to splenic bacterial accumulation, we performed targeted RT-qPCR in the spleen across three functional modules: the cGAS-STING-IFN axis (**Fig. 3A**), pro-inflammatory effectors (**Fig. 3B**), and Hv1 redox-related genes (**Fig. 3C**). Within the cGAS-STING axis, Hv1KO/TBI spleens showed coordinated upregulation of the cytosolic DNA sensors *Mb21d1* (cGAS) and *Tmem173* (STING), together with the interferon-stimulated GTPase gene *Irgm2*, relative to both Hv1KO/Sham and WT/TBI; *Irf3* was elevated in Hv1KO/TBI by simple-effects comparison (**Fig. 3A**). In contrast, the canonical type I IFN output *Ifnb1* was suppressed in Hv1KO/TBI relative to both Hv1KO/Sham and WT/TBI. Among pro-inflammatory effectors, *Fcgr4*, *Tnfrsf1b*, *Il17ra*, *Jak3* were significantly elevated specifically in Hv1KO/TBI spleens with significant interaction and injury effects, and *Syk* showed a similar trend (**Fig. 3B**). Despite *Hv1* loss, *Cybb* (NOX2), *Id2* and *Prkca* were significantly elevated in Hv1KO/TBI spleen (**Fig. 3C**). Together, the Hv1-deficient spleen sustains a chronic immune active state characterized by engaged DNA sensing, suppressed IFN-β output, elevated JAK- and Fc receptor-associated inflammatory signaling, and increased NADPH oxidase-related expression, a molecular signature that coincides with splenic 16S accumulation and suppression of antibacterial effector pathways, identifying the Hv1KO spleen as a site of chronic inflammation decoupled from effective IFN-β-dependent antibacterial defense.

**Fig. 3.**
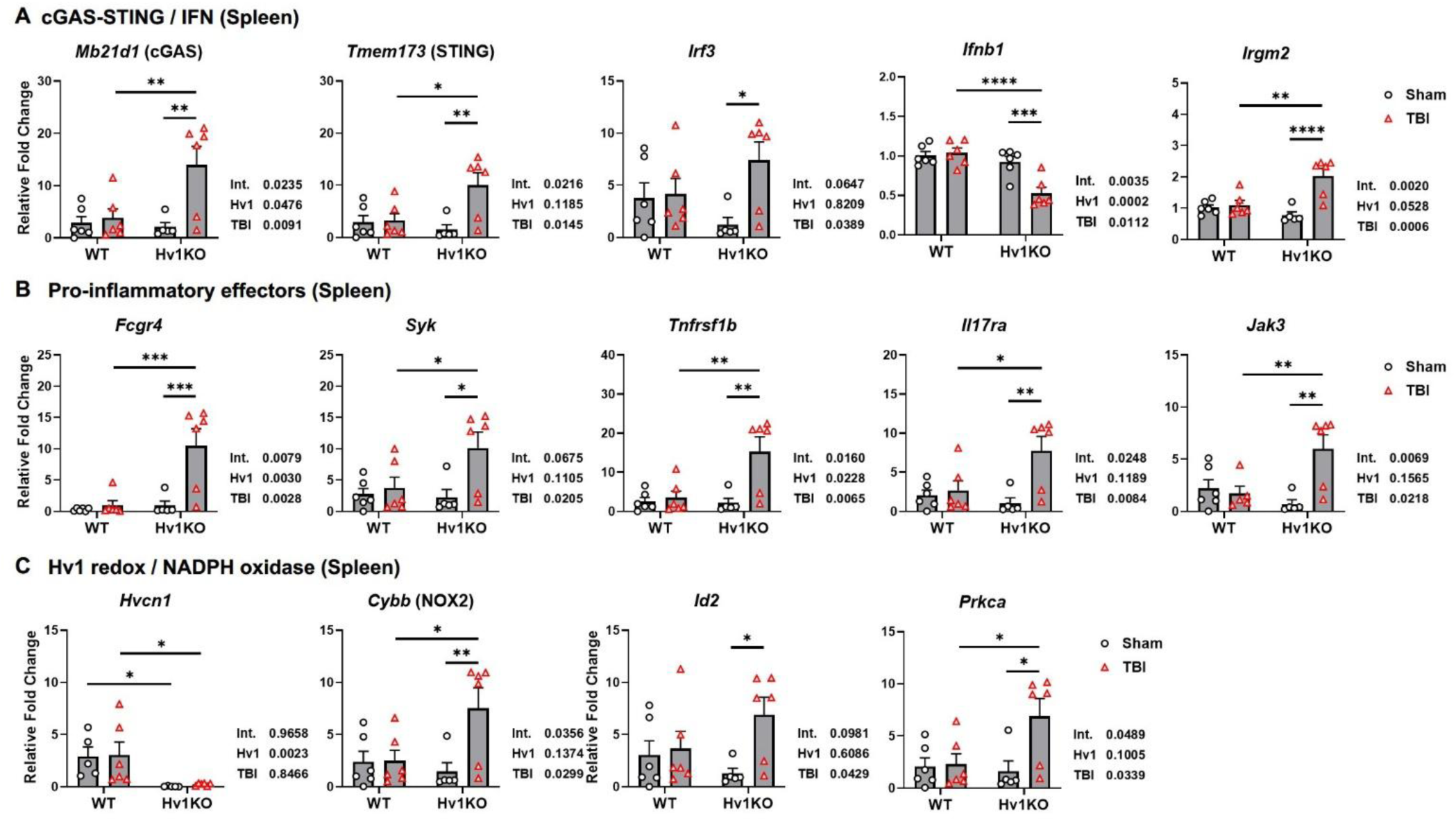
Splenic cGAS-STING-IFN signaling and pro-inflammatory effectors are dysregulated in Hv1KO mice after chronic TBI. **(A)** Hv1KO/TBI spleens showed elevated *Mb21d1*, *Tmem173*, and *Irgm2* but reduced *Ifnb1* relative to WT/TBI and Hv1KO/Sham; increased *Irf3* in Hv1KO/TBI compared with Hv1KO/Sham. **(B)** Pro-inflammatory effector genes *Fcgr4*, *Syk*, *Tnfrsf1b*, *Il17ra*, and *Jak3* were elevated in Hv1KO/TBI spleen. **(C)** *Hvcn1* mRNA was absent in both Hv1KO groups; *Cybb* (NOX2), *Id2*, and *Prkca* were elevated in Hv1KO/TBI spleen. Sample sizes: n=5-6 mice/group. Statistical analyses: Two-way ANOVA with Šídák-corrected simple-effects comparisons (A–C) with p-values representing interaction (Int.), *Hvcn1* knock-out (Hv1), and TBI main-effect shown beside each panel. *p≤0.05, **p≤0.01, ***p≤0.001, ****p≤0.0001.

### Hv1 loss impairs splenic neutrophil phagocytosis and dysregulates the oxidative burst under chronic TBI

The transcriptomic collapse of pathogen-defense programs in the Hv1KO/TBI spleen raised the question of whether splenic neutrophils were functionally compromised. Specifically, we gated out splenic neutrophils and measured particle uptake, lysosome activation, and ROS production by flow cytometry (**Fig. 4**). The phagocytic capacity of pHrodo-labeled *S. aureus* bioparticles in splenic neutrophils differed by genotype but not by injury, and was significantly reduced in Hv1KO/TBI compared to WT/TBI (**Fig. 4A**). Engulfment of pHrodo-labeled *E. coli* bioparticles showed a stronger genotype effect, with significant reductions in Hv1KO relative to WT under both sham and injured conditions (**Fig. 4B**). Engulfment of 0.5-µm latex beads and Fc receptor-dependent IgG beads carried the same genotype dependence and similar pattern between the two injured groups (**Fig. 4C-D**). Furthermore, CD68 expression on splenic neutrophils declined after chronic TBI, and the reduction was significant within the Hv1KO groups (**Fig. 4E**). DHR123 signal showed significant genotype, injury, and interaction effects: it was highest in Hv1KO/Sham neutrophils, fell back to WT level post-injury, unchanged across the two WT groups (**Fig. 4F**).

**Fig. 4.**
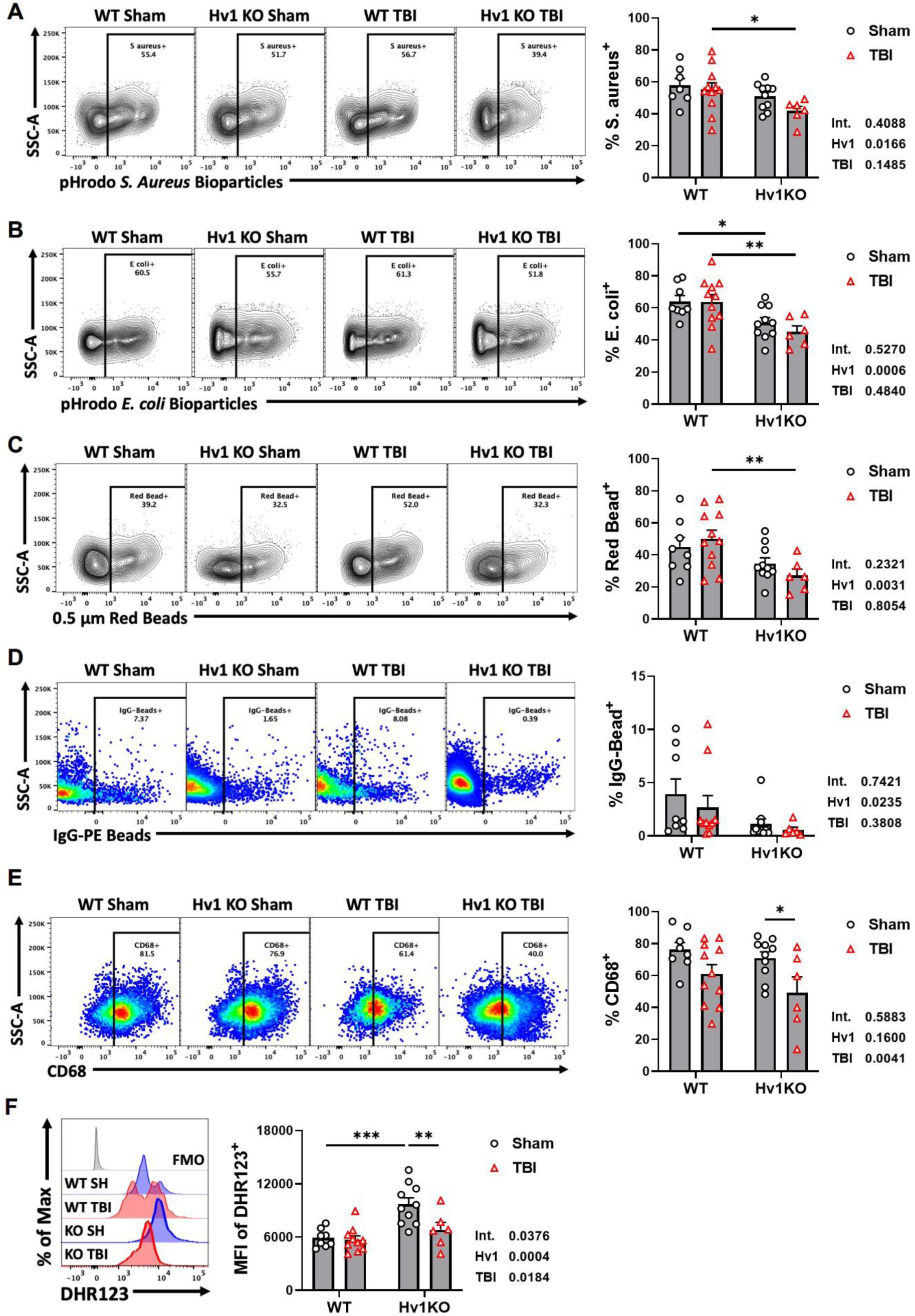
Hv1 deficiency impairs splenic neutrophil phagocytosis and dysregulates the oxidative burst under chronic TBI. **(A)** The uptake of pHrodo-labeled S. Aureus bacterial particles by splenic neutrophil was altered with notable genotype effect and significantly impaired in Hv1KO/TBI compared with WT/TBI. **(B)** Splenic neutrophil uptake of pHrodo-labeled E. coli bacterial particles was significantly reduced by Hv1 deficiency in both paired comparisons respectively. **(C)** Hv1 deficiency significantly impaired neutrophil phagocytic capacity of a broader bead-engulfment after chronic TBI. **(D)** Hv1 deficiency showed notable genotype effect in Fc-receptor-dependent phagocytosis of splenic neutrophils without significant differences between groups. **(E)** CD68 expression was reduced in splenic neutrophils in post-injury spleen compared with Sham, which showed significant difference between two Hv1KO groups. **(F)** DHR123 signal reached the highest level in Hv1KO/Sham splenic neutrophils compared with WT/Sham and Hv1KO/TBI, showing significant effects of genotype and injury as well as their interaction. Sample sizes: WT/Sham, n=7-8; WT/TBI, n=10-11; Hv1KO/Sham, n=10, Hv1KO/TBI, n=6. Statistical analyses: Two-way ANOVA with Šídák-corrected simple-effects comparisons (A-F) with p-values representing interaction (Int.), *Hvcn1* knock-out (Hv1), and TBI main-effect shown beside each panel. *p≤0.05, **p≤0.01, ***p≤0.001, ****p≤0.0001.

Across these assays, Hv1 deficiency is the principal determinant of phagocytic uptake, and the impairment is most pronounced where genotype and injury converge. The lysosome marker CD68 falls predominantly with injury in Hv1 deficient neutrophils, suggesting impaired phagolysosomal capacity. Hv1 loss also changed neutrophil ROS constitutively with altered responses to chronic TBI.

### Hv1 deficiency imposes a constitutive innate immune activation in the lung, amplified by chronic TBI

Having established that Hv1 deficiency drives spleen-restricted antibacterial failure and chronic inflammatory decompensation, we asked whether the lung, the primary site of bacterial clearance for inhaled pathogens, followed the same trajectory. Lung profiling with the same NanoString panel revealed a fundamentally different statistical architecture from the spleen. Hierarchical clustering and PLS-DA (39.5% variance) separated all four groups (**Fig. 5A-B**). ssGSEA revealed a predominantly genotype-driven immune bias in Hv1KO lungs (**Fig. 5C**, **supplemental Table 3**). Eight pathways were suppressed in Hv1KO lung irrespective of injury, including Angiogenesis, Fc Receptor and Cytokine Signaling, Cell Cycle and Apoptosis, T-cell Activation and Checkpoint Signaling, Growth Factor Signaling, Differentiation and Maintenance of Myeloid Cells, and Metabolism. Seven pathways were elevated in the opposite direction, including Antigen Presentation, Lymphocyte activation, Pathogen Response, Chemokine, Complement, and TLR signaling, TH1 Activation. Additionally, Cell Migration and Adhesion, and TH2 Activation, were marked chiefly by suppression in WT/TBI; ECM remodeling and Interferon Signaling, on the other hand, by elevation in Hv1KO/Sham. Two-way ANOVA confirmed significant genotype effects for most pathways, with only Complement Activation showing an injury main effect and no significant gene *x* injury interactions.

**Fig. 5.**
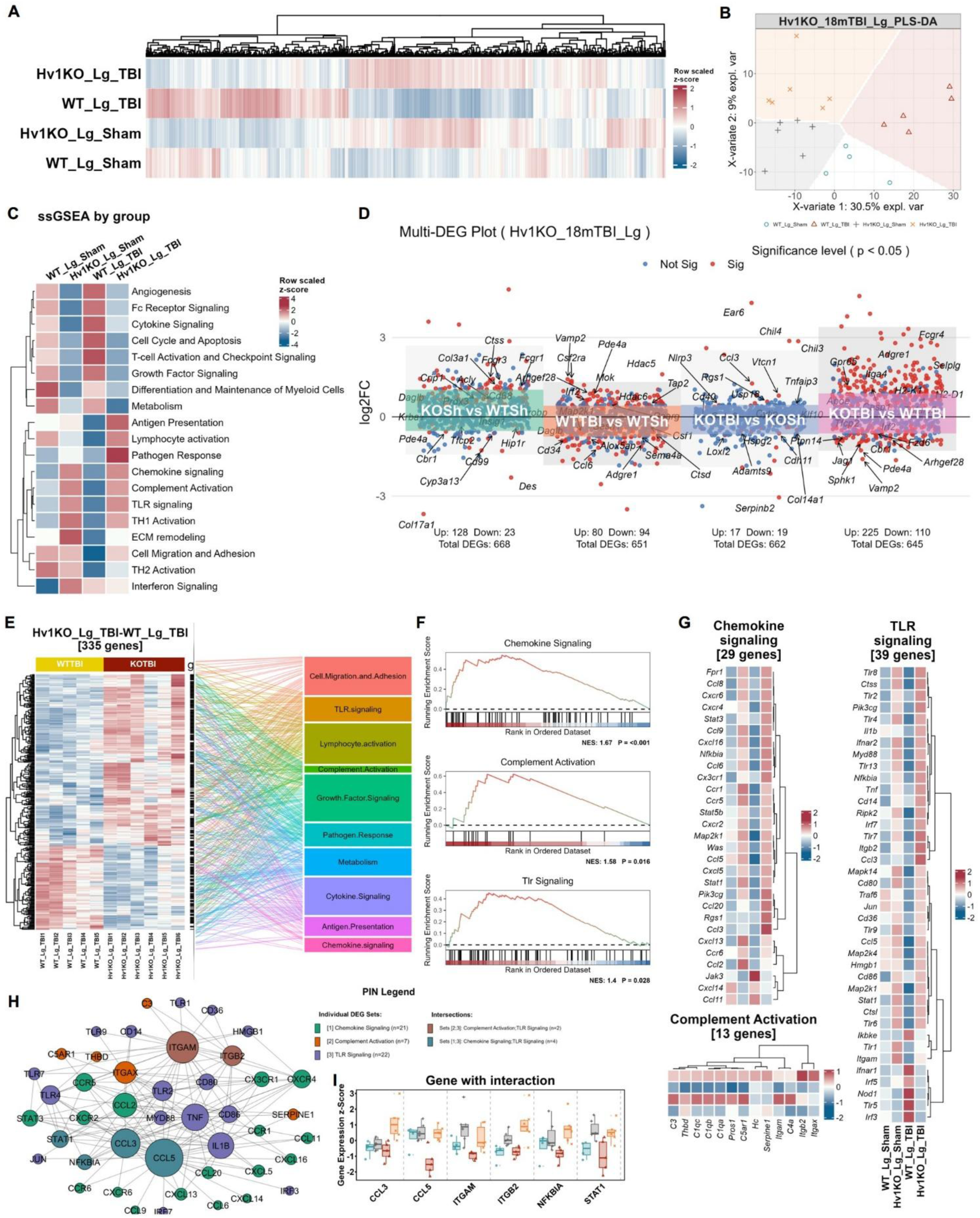
Hv1 loss enhanced immune responses after chronic TBI in the lung. **(A)** Myeloid Innate Immunity panel generates distinct gene heatmaps from the lung (Lg) tissue of the four groups. **(B)** PLS-DA plot indicates different transcriptomic signatures by separated group clusters. **(C)** The heatmap from single-sample Gene Set Enrichment Analysis (ssGSEA) of Myeloid Innate Immunity panel highlights predominant differences caused by Hv1 deficiency. **(D)** DEGs with Hv1 KO versus WT or TBI versus Sham were demonstrated in the multi-volcano graph. **(E)** DEGs with Hv1 KO TBI versus WT TBI in the lung mostly cluster into the top 10 enriched pathways. **(F)** Customized GSEA analysis demonstrates significant activation of “Chemokine Signaling”, “Complement Activation”, and “Tlr Signaling” based on DEGs from (E). **(G)** Gene expression profiles of DEGs from the three activated pathways (G) are exhibited as z-score heatmaps. **(H)** PIN demonstrates theoretical interactions between the DEGs derived proteins. **(I)** Expression profiles across groups of the six genes with inter-pathway interactions. Sample sizes: WT/Sham, n=4; Hv1KO/Sham, n=6; WT/TBI, n=5; Hv1KO/TBI, n =6.

Furthermore, between-group differential expression analysis (**Fig. 5D**) revealed that the largest DEG set arose specifically from the comparison of Hv1KO/TBI vs WT/TBI: 335 DEGs (225 up, 110 down), exceeding the genotype effect in sham animals (151 DEGs) and the injury effect within either genotype (174 in WT, 36 in Hv1KO). These DEGs derived from Hv1 deficiency with injury clustered into ten top-ranked pathways, prominently Cell Migration and Adhesion, TLR Signaling, Lymphocyte Activation, Complement Activation, Growth Factor Signaling, and Pathogen Response (**Fig. 5E**). Panel-based GSEA confirmed activation of Chemokine Signaling, Complement Activation, and TLR Signaling (**Fig. 5F**), with coordinated upregulation of chemokines and receptors (*Cxcr4*, *Cxcl16*, *Ccl9*, *Ccl5*, *Ccl3*, *Ccl11*, *Cx3cr1*), TLR components (*Tlr8*, *Tlr2*, *Tlr13*, *Ctss*, *Il1b*, *Ifnar2*, *Tnf*, *Irf7*), and complement components (*C3*, *C1qa*/*b*, *Hc*, *Itgam*, *Itgax*) (**Fig. 5G**). PIN analysis identified CCL3, CCL5, ITGAM, ITGB2, and STAT1 as central hubs with extensive cross-pathway connectivity (**Fig. 5H**); each was specifically elevated in Hv1KO/TBI lungs relative to WT/TBI (**Fig. 5I**), defining a coordinated module driving the pulmonary response.

Additionally, factor-decomposition analyses dissected gene-level contributions respectively (**Supplemental Fig. 3A**). The gene *x* injury interaction identified 63 DEGs with significant GSEA activation of Pathogen Response (**Supplemental Fig. 3B**), opposite from the spleen. The genotype effect produced 131 DEGs with GSEA activation of Chemokine Signaling and TLR Signaling (**Supplemental Fig. 3C**), reinforcing the constitutive bias observed at the pathway-score level. The injury effect produced 130 DEGs with GSEA suppression of Cell Migration and Adhesion (**Supplemental Fig. 3D**). Meanwhile, we examined the lung in three functional modules by qPCR. In the cGAS-STING-IFN axis (**Supplemental Fig. 4A**), *Mb21d1*, *Tmem173*, and *Irf3* were higher in Hv1KO than WT lungs. *Ifnb1* showed a genotype-dependent injury response, rising with TBI in WT but declined in Hv1KO, and *Ccl5* was elevated by both Hv1 deletion and injury, peaking in Hv1KO/TBI. In the TLR signaling and inflammatory module (**Supplemental Fig. 4B**), *Tlr13*, *Tlr8*, and *Clec7a* were upregulated in Hv1KO lungs with significant genotype effect, consistent with the NanoString genotype contrast (**Supplemental Fig. 3C**). *Ctss* increased with both genotype and injury, and *Il1b* was higher in Hv1KO groups. In the Hv1-NOX2 and myeloid effector module (**Supplemental Fig. 4C**), *Hvcn1* mRNA was absent in Hv1KO lungs, confirming the knockout, while its functional partner *Cybb* (NOX2) was elevated with Hv1 loss. *Itga4* and *Fcgr4* were both higher in Hv1KO groups, and *Batf3* showed a significant interaction effect, increasing with Hv1 deficiency to a comparable level with TBI in WT.

Collectively, unlike the injury-driven decompensation observed in the spleen, the lung exhibited a constitutively biased, genotype-dominated innate immune response. Hv1 deficiency alone reprogrammed the pulmonary transcriptome toward enhanced chemokine, TLR, and complement signaling at both the pathway and gene levels, with chronic TBI further amplifying these responses.

### Hv1 loss remodels hepatic immune-metabolic signaling with chronic TBI

With these organ-divergent responses in spleen and lung, we turned to the liver, a blood-filtering organ and central hub of systemic bacterial clearance, to investigate which trajectory its immune compartment would follow through the same NanoString panel. Hierarchical clustering demonstrated distinct transcriptomic profiles by group, especially in Hv1KO/TBI liver, with PLS-DA separated roughly different groups (**Fig. 6A-B**). In contrast to the spleen and lung, ssGSEA in the liver revealed an interaction-dominant architecture in which Hv1 deficiency and TBI converged to reshape pathway activities (**Fig. 6C, Supplemental Table 4**). Pathways linked to immune effector and antibacterial functions (Chemokine Signaling, Complement Activation, Pathogen Response, Interferon and Cytokine Signaling, Antigen Presentation) were relatively muted in WT/TBI but strongly activated in Hv1KO/TBI, while metabolic, trophic, and lymphocyte programs (Metabolism, Growth Factor and Fc Receptor Signaling, Angiogenesis, Lymphocyte Activation) were oppositely boosted in WT/TBI yet suppressed in Hv1KO/TBI. This bidirectional partition contrasts with the spleen, where Hv1KO baseline activation decompensates under TBI, and with the lung, where Hv1 imposes a largely constitutive inflammatory bias, defining the liver as a synergy-driven mode of Hv1-dependent remodeling.

**Fig. 6.**
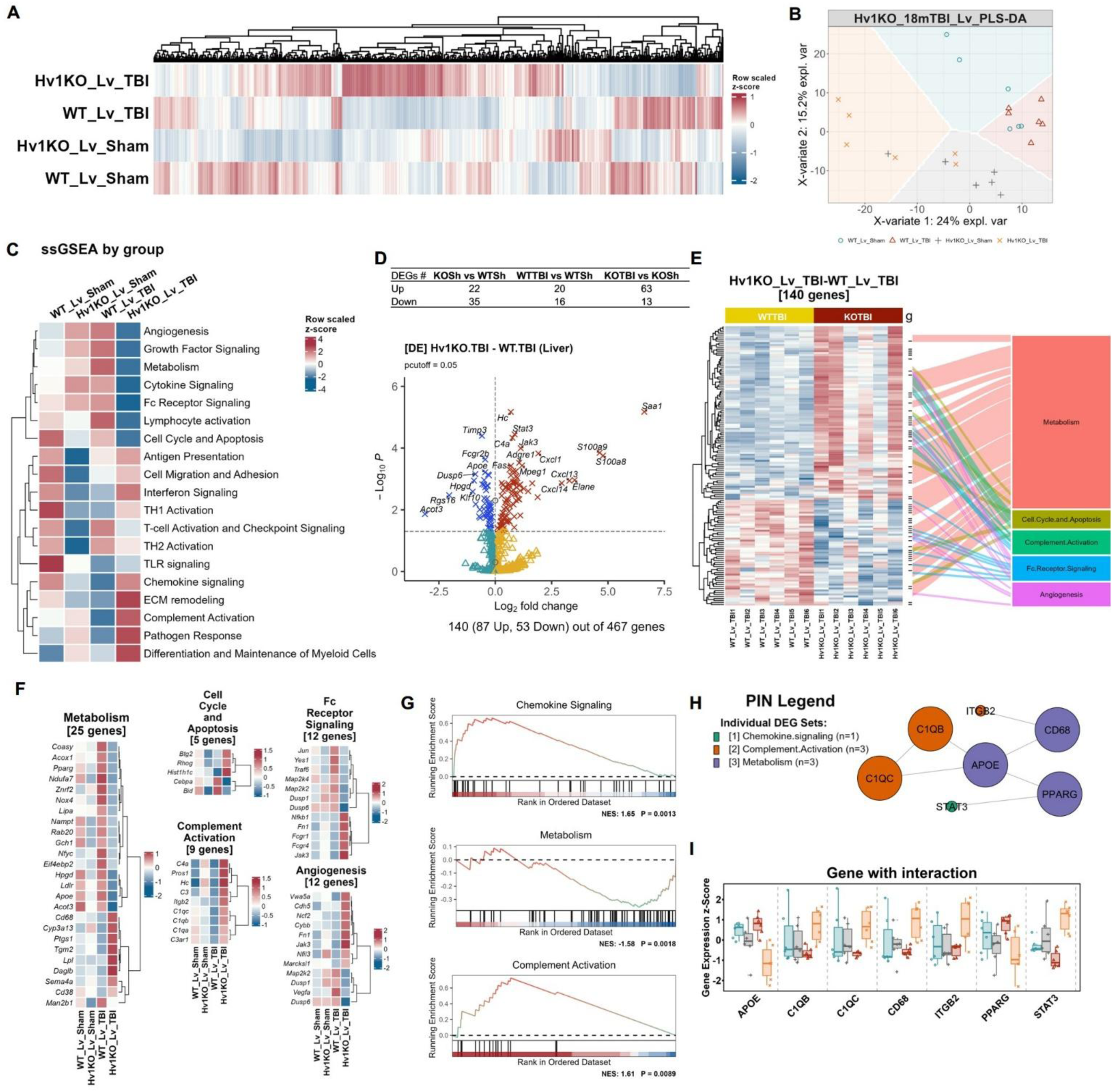
Hv1 loss with chronic TBI convergently dysregulated immune responses and metabolism signaling in the liver. **(A)** Myeloid Innate Immunity panel generates distinct gene heatmaps from the liver (Lv) tissue of the four groups. **(B)** PLS-DA plot indicates different transcriptomic signatures by separated group clusters. **(C)** The heatmap from ssGSEA of Myeloid Innate Immunity panel exhibits distinct post-injury alterations caused by Hv1 deficiency. **(D)** The volcano graph displays DEGs in Hv1 KO liver versus WT. **(E)** The heatmap demonstrates gene expression profile of the significantly changed DEGs, most of which cluster into the top 5 pathways. **(F)** Expression heatmaps of DEGs between Hv1 KO TBI versus WT TBI in the top 5 pathways, showing across-group comparisons. **(G)** GSEA analysis indicates significant suppression of “Metabolism”, as well as activation of “Chemokine Signaling” and “Complement Activation”. **(H-I)** PIN from the 3 pathways (G) implies potential interactions based on DEGs with expression displayed (I). Sample sizes: WT/Sham, n=6; Hv1KO/Sham, n=6; WT/TBI, n=6; Hv1KO/TBI, n=6.

The direct comparison of Hv1KO/TBI vs WT/TBI generated 140 DEGs, substantially more than either the baseline genotype effect (57 DEGs) or the injury effect within either genotype (36 in WT, 76 in Hv1KO) (**Fig. 6D**). The 140 DEGs are mostly clustered into Metabolism, Cell Cycle and Apoptosis, Complement Activation, Fc Receptor Signaling, and Angiogenesis (**Fig. 6E-F**), with coordinated upregulation of inflammatory and complement effectors (*C1qa*/*b*/*c*, *Fn1*, *Fcgr1*, *Jak3*) and downregulation of metabolic-related genes (*Acot3*, *Apoe*, *Hpgd*, *Pparg*). Panel-based GSEA on the KO/TBI vs WT/TBI comparison confirmed activation of Chemokine Signaling and Complement Activation alongside suppression of Metabolism (**Fig. 6G**). PIN analysis identified potential connections between metabolic and immune nodes, including metabolism (APOE, CD68, PPARG), complement components (C1QB, C1QC, ITGB2), and chemokine signaling (STAT3) (**Fig. 6H**). APOE and PPARG were reduced in Hv1KO/TBI alongside coordinated augments of others in connecting nodes (**Fig. 6I**). Moreover, factor-decomposition analyses revealed the interaction-dominant hepatic phenotype (**Supplemental Fig. 5**). The genotype effect induced 121 DEGs (19 up, 102 down) with GSEA suppression of Chemokine Signaling and T-cell Activation (**Supplemental Fig. 5A-B**). The injury effect produced 41 DEGs (10 up, 31 down) with GSEA suppression of Chemokine Signaling, TLR Signaling, Pathogen Response, and ECM Remodeling (**Supplemental Fig. 5C**). The gene *x* injury interaction identified 167 DEGs (140 up, 27 down) with significant GSEA activation of Chemokine Signaling, Pathogen Response, Complement Activation, and TLR Signaling (**Supplemental Fig. 5D**), pathways suppressed by either factor alone. Thus, the hepatic phenotype is interaction-dominant, coupling complement and inflammatory activation to metabolic suppression specifically where Hv1 deficiency meets chronic TBI.

To validate the NanoString findings at single-gene resolution, RT-qPCR was performed in the liver covering four functional modules. (1) cGAS-STING axis (**Supplemental Fig. 6A**): *Mb21d1* (cGAS), *Tmem173* (STING), and *Ifnb1* showed no significant changes; the cytokine-signaling kinase *Jak3* was elevated by Hv1 deficiency, reaching significance in Hv1KO/TBI versus WT/TBI. (2) Oxidative stress module (**Supplemental Fig. 6B**): Hvcn1 ablation in both Hv1KO groups confirmed the constitutive knockout; *Cybb* (NOX2) and *Hmox1* were unchanged. (3) Lipid metabolism(**Supplemental Fig. 6C**): *Acot3* was reduced by both genotype and injury, with significant decrease by TBI within WT liver, and *Hpgd* was reduced by Hv1 deficiency; in contrast, *Cyp3a13* was significantly higher in Hv1KO/TBI than WT/TBI, and *Apoe* responded only to injury, with no genotype or interaction effect. (4) Tissue remodeling / repair signaling (**Supplemental Fig. 6D**): *Dusp6* exhibited significant interaction and injury effect, with TBI-driven significant suppression in Hv1KO liver; *Tgm2* was elevated by Hv1 deficiency, reaching significance in Hv1KO/TBI versus WT/TBI; *Hgf* showed a main effect of injury, lower after TBI in both genotypes.

Together, the liver presents a third organ-specific phenotype characterized by interaction-dominant remodeling. The pulmonary and hepatic transcriptional phenotypes contrast sharply with the spleen without bacterial burden. Hv1 loss therefore produces organ-specific patterns of chronic TBI immune dysregulation rather than a generalized peripheral response.

### Hv1 deficiency reshapes peripheral immune cell composition after chronic TBI in an organ-specific manner

To dissect the leukocyte subpopulations implementing the organ-specific pathway signatures, we deconvolved the NanoString profiles into immune cell-type enrichment and validated key shifts by flow cytometry in spleen, lung, and liver. The enrichment scores of each cell type were put together by group as scaled z-score heatmap, and cell types with a significant genotype, injury, or interaction term were shown as box plots.

In the spleen (**Fig. 7A–B; Supplemental Table 5A**), neutrophil and macrophage scores exhibited significant genotype × injury interactions and were highest in the Hv1KO/Sham group. Mast cell and exhausted CD8 T-cell scores were increased in Hv1KO mice independent of injury, whereas B-cell scores were reduced in Hv1KO. In contrast, the CD45 signal was selectively decreased in Hv1KO/Sham, also exhibiting a significant genotype × injury interaction. By flow cytometry, CD11b⁺ and Ly6G⁺ cells were both highest in Hv1KO/TBI spleen, each elevated significantly relative to WT/TBI and to Hv1KO/Sham (**Fig. 7C**). In the lung (**Fig. 7D-E; Supplemental Table 5B**), cytotoxic T-cell scores were comparable at baseline but decreased in WT while remaining preserved in Hv1KO following TBI, resulting in significant genotype, injury, and genotype × injury interaction effects. For all other immune cell types and the CD45 marker, genotype was the predominant driver, with consistently higher levels in Hv1KO than WT. Flow cytometry corroborated this constitutive myeloid bias, demonstrating significantly increased frequencies of CD11b⁺ and Ly6G⁺ cells in Hv1KO lungs at baseline (**Fig. 7F**). In the liver (**Fig. 7G–H; Supplemental Table 5C**), neutrophil and macrophage scores were more strongly enriched in Hv1KO than WT following TBI, with significant genotype × injury interaction effects. CD45 expression followed a similar pattern. Flow cytometry validated these findings, revealing a significant increase in CD11b⁺ cells in Hv1KO/TBI livers compared with WT/TBI, along with significantly elevated Ly6G⁺ neutrophils driven by both Hv1 deficiency and chronic TBI (**Fig. 7I**). Across all three organs, immune cell composition was remodeled by Hv1 loss after chronic TBI in an organ-specific way, while expansion of CD11b⁺ myeloid cells and Ly6G⁺ neutrophils emerged as a shared Hv1-dependent axis. We interpret enrichment scores as transcriptional state and flow counts as cell population abundance, two complementary measures targeting different aspects.

**Fig. 7.**
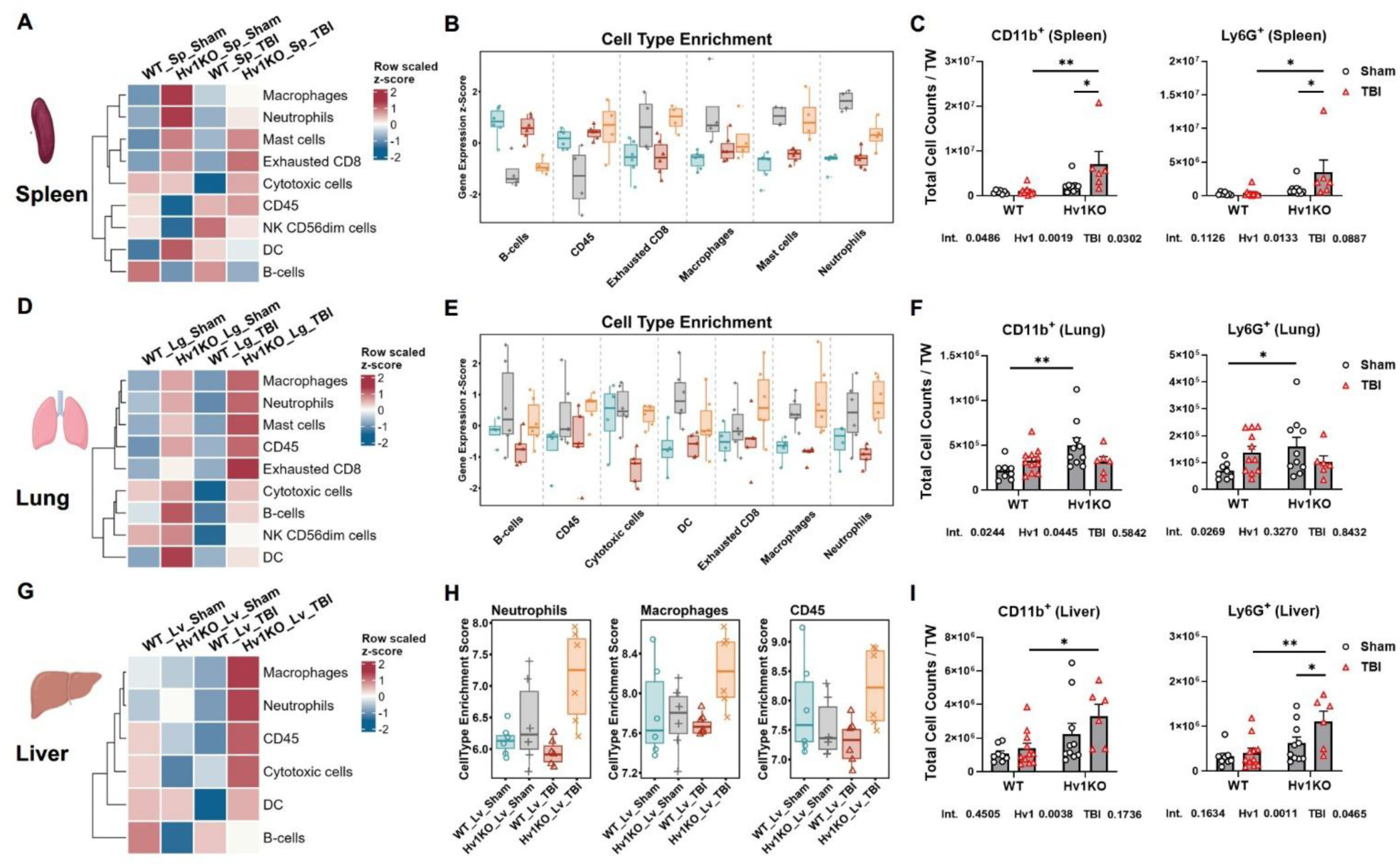
Hv1 deficiency reshapes peripheral immune cell composition in an organ-specific manner after chronic TBI. Cell-type enrichment was deconvolved from NanoString Myeloid Innate Immunity panel data and validated by flow cytometry in spleen, lung, and liver. **(A)** Clustered heatmap of cell-type enrichment scores in spleen (row-scaled z-score). **(B)** Box plots of spleen cell types with a significant genotype, injury, or interaction term. **(C)** Flow cytometry of total CD11b⁺ (myeloid) and Ly6G⁺ (neutrophil) splenocytes, expressed as total cell counts normalized by tissue weight (TW). **(D-E)** Enrichment-score heatmap (D) and box plots (E) for lung. **(F)** Flow cytometry of CD11b⁺ and Ly6G⁺ lung cells. **(G-H)** Enrichment-score heatmap (G) and box plots (H) for liver. **(I)** Flow cytometry of CD11b⁺ and Ly6G⁺ liver cells. Sample sizes: n=5-6 mice/group. Statistical analyses: Two-way ANOVA with Šídák-corrected simple-effects comparisons (C, F, I) with p-values representing interaction (Int.), *Hvcn1* knock-out (Hv1), and TBI main-effect shown beside each panel. *p≤0.05, **p≤0.01, ***p≤0.001, ****p≤0.0001.

### Loss of Hv1 impairs gut barrier and mucus integrity after chronic TBI

The spleen-restricted accumulation of pan-bacterial 16S rRNA in Hv1KO/TBI mice, and the absence of classical opportunistic pathogens *E. coli* and *S. aureus*, along with the suppression of splenic antibacterial pathways under chronic injury, pointed to commensal bacterial translocation as the most likely source. Because the intestinal epithelium is the primary interface separating commensal bacteria from systemic circulation [3], we assessed colonic barrier, mucus, redox/sensing, and effector genes by RT-qPCR across three modules: epithelial barrier and mucus defense (**Fig. 8A**), cGAS-STING-IFN and Hv1-NOX2 redox (**Fig. 8B**), and inflammatory and anti-microbial effectors (**Fig. 8C**).

**Fig. 8.**
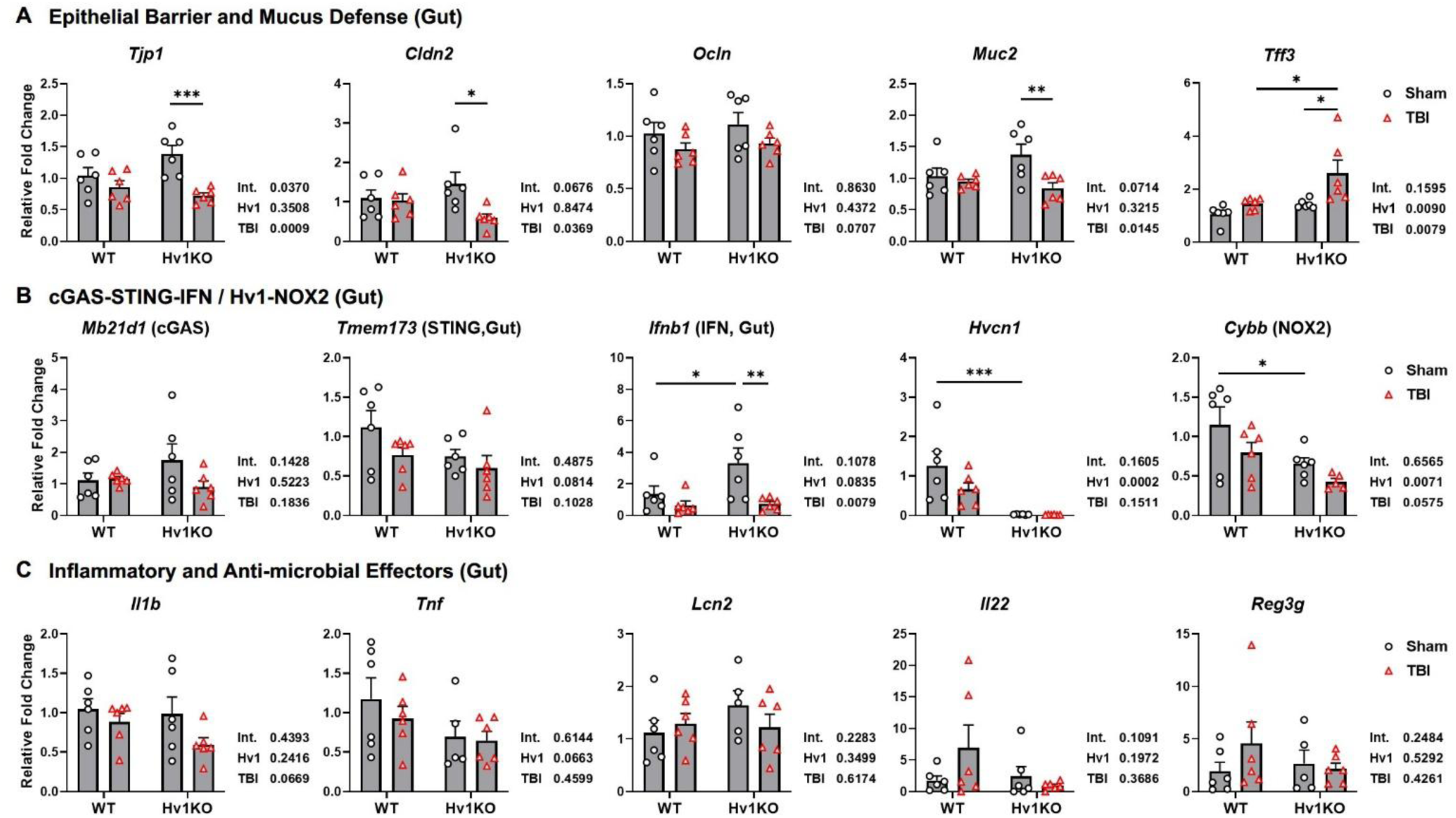
Loss of Hv1 compromises colonic barrier integrity and mucosal innate defense after chronic TBI. Colonic RT-qPCR across three functional modules in all four groups. **(A)** Epithelial barrier and mucus defense: *Tjp1* (ZO-1), *Cldn2* (claudin-2), *Ocln* (occludin), *Muc2*, *Tff3*. **(B)** cGAS-STING-IFN sensing and Hv1-NOX2 redox: *Mb21d1* (cGAS), *Tmem173* (STING), *Ifnb1*, *Hvcn1*, *Cybb* (NOX2). **(C)** Inflammatory and antimicrobial effectors: *Il1b*, *Tnf*, *Lcn2*, *Il22*, *Reg3g*. Sample sizes: n=5-6 mice/group. Statistical analyses: two-way ANOVA with Šídák-corrected simple-effects comparisons; interaction (Int.), Hv1 (*Hvcn1* knockout), and TBI main-effect p-values are shown beside each panel. *p≤0.05, **p≤0.01, ***p≤0.001, ****p≤0.0001.

Among barrier and mucus genes (**Fig. 8A**): *Tjp1* (ZO-1), the master scaffolding protein of the tight junction complex, showed a significant gene *x* injury interaction and injury effect, which was modestly elevated in Hv1KO/Sham relative to WT/Sham but dramatically suppressed after TBI. *Cldn2* (Claudin-2), a tight junction channel protein highly expressed in intestinal crypts, was significantly suppressed specifically in Hv1KO/TBI with significant injury effect. *Ocln* (Occludin), a key protein maintaining tight junction integrity for macromolecule-selective barrier, showed a marginal decline trend post-injury. *Muc2* (Muc2), the major secretory mucin and primary physical barrier between the lumen and the epithelium, showed significant TBI-driven suppression specifically in Hv1KO with significant injury effect and marginal interaction. *Tff3* (trefoil factor 3), an epithelial repair factor, was significantly elevated in Hv1KO/TBI versus both Hv1KO/Sham and WT/TBI with significant genotype and injury effects. No significant differences were detected between two WT groups in this module, implying that Hv1 deficiency hindered post-injury recovery of gut barrier and mucus integrity. Within the redox and DNA-sensing module (**Fig. 8B**): *Mb21d1* (cGAS) and *Tmem173* (STING) were unchanged across groups. *Ifnb1* peaked in Hv1KO/Sham and declined in Hv1KO/TBI with a significant injury effect. *Hvcn1* mRNA was absent in both Hv1KO groups, confirming the colonic knockout. *Cybb* (NOX2) was reduced in Hv1KO intestine with a significant genotype effect and a marginal injury trend. Target effector gene expression showed no significant changes across groups (**Fig. 8C**), including *Il1b*, *Tnf*, and *Lcn2*, as well as the antimicrobial *Il22* and *Reg3g*.

Therefore, the colonic data identify a coordinated defect in Hv1KO/TBI tissue spanning three layers of mucosal protection: (1) loss of the *Tjp1* tight-junction scaffold, (2) suppression of the *Muc2* mucus barrier with a compensatory but insufficient *Tff3* repair response, and (3) reduced NOX2-dependent ROS capacity and type I IFN output, uncompensated by the IL-22–REG3 antimicrobial program. Their concurrence provides a potential molecular framework for commensal bacterial translocation underlying the spleen-restricted 16S accumulation.

### Hv1 loss confers modest behavioral resilience with altered cortical transcriptional responses after chronic injury

We next assessed cognitive, social, and affective function across all four groups (Schemed as **Supplemental Fig. 7A**). Social behavior was assessed using the three-chamber Social Recognition (SR) test. During habituation, no group showed significant cup preference (**Supplemental Fig. 7B**), confirming no innate bias. In the social preference phase (**Supplemental Fig. 7C**), all groups showed preference for the stranger-containing chamber, with no significant effects of genotype, injury, or interaction. In the social novelty preference phase, a robust gene *x* injury interaction and significant main effects were observed (**Supplemental Fig. 7D**). WT/TBI mice showed significantly impaired social novelty preference compared with WT/Sham, reflecting a chronic TBI-induced deficit in social recognition memory; however, Hv1KO/TBI mice retained social novelty preference at levels comparable to Sham groups and significantly outperformed WT/TBI mice, indicating that Hv1 deficiency preserves social recognition after chronic TBI. Affective behavior was assessed using the Novelty-Suppressed Feeding (NSF) test, where prolonged latency to feed in a novel arena reflects anxiety- or depressive-like behavior. WT/TBI mice showed significantly increased latency relative to WT/Sham (**Supplemental Fig. 7E**), reflecting chronic TBI-induced affective dysfunction; meanwhile, Hv1KO/TBI mice showed reduced latency relative to WT/TBI, indicating partial rescue. NSF latency in the home cage showed no significant differences across groups (**Supplemental Fig. 7F**), confirming that the novel-arena finding reflects affective rather than locomotor or motivational deficits. Cognitive functions were assessed using the Y-maze, in which spontaneous alternation, a measure of spatial working memory, reduced by TBI in WT but retained in Hv1KO with marginal interaction trends (**Supplemental Fig. 7G**). Hv1KO mice showed significantly less total arm entries than WT counterparts (**Supplemental Fig. 7H**), indicating decreased exploratory activity with Hv1 deficiency. Recognition memory was assessed using the Novel Object Recognition (NOR) test. Baseline object bias was equivalent across groups (**Supplemental Fig. 7I**), and novelty preference showed a significant injury effect with reduced novelty exploration comparable in two TBI groups (**Supplemental Fig. 7J**). Open field analysis showed no significant differences in total distance traveled or maximum speed (**Supplemental Fig. 7K-L**), confirming that the social and affective phenotypes were not confounded by gross locomotor abnormalities.

To relate these behavioral phenotypes to cortical molecular changes, we profiled ipsilateral somatosensory cortex using NanoString Neuropathology panel. Hierarchical clustering and PLS-DA separated all four groups along genotype- and injury-driven axes (**Supplemental Fig. 8A-B**). Hv1 deficiency and chronic TBI jointly reshape a limited set of neuropathology pathways rather than globally rewiring the panel. Two-way ANOVA on ssGSEA scores identified Angiogenesis, Cytokines, and Tissue Integrity as the only pathways with significant gene *x* injury interactions, with Angiogenesis and Cytokines showing the highest scores and Tissue Integrity the lowest specifically in Hv1KO/TBI cortex (**Supplemental Fig. 8C; Supplemental Table 6**). Several other pathways displayed main effects without interaction. At the genotype level, Hv1 loss modestly activated Angiogenesis, Cytokines, Lipid Metabolism, and Oxidative Stress and suppressed Tissue Integrity, whereas TBI alone enhanced Angiogenesis, Cytokines, Matrix Remodeling, Neuronal Cytoskeleton, and Tissue Integrity while impaired Growth Factor Signaling, Lipid Metabolism, Oxidative Stress, and Apoptosis. The resulting ssGSEA heatmap therefore depicts a cortical phenotype in which chronic TBI in Hv1-deficient mice amplifies inflammatory and vascular remodeling programs while further eroding tissue-structural pathways. Differential expression of Hv1KO/TBI versus WT/TBI yielded 103 DEGs overlapping with other comparisons (**Supplemental Fig. 8D; Supplemental Fig. 9A**), which demonstrated upregulation of immediate-early and stress-response transcripts (*Fos*, *Xbp1*, *Atf4*, *Ptgs2*), Cxcl12, the TWEAK receptor *Tnfrsf12a*, and the angiogenic regulator *Egfl7*, together with downregulation of neuronal connectivity genes (*Cntn1*, *Nrxn1*) and the astroglial water channel *Aqp4*. The DEG set mostly clustered into the top 5 pathways of Oxidative Stress, Transcription and Splicing, Unfolded Protein Response, Carbohydrate Metabolism, and Autophagy (**Supplemental Fig. 8E**). Panel-based GSEA confirmed activation of Oxidative Stress and Cytokines with suppression of Tissue Integrity and Myelination (**Supplemental Fig. 8F**). Cross-group gene expression profiles were displayed for the DEGs significantly altered by Hv1 deficiency after TBI (**Supplemental Fig. 8G**). PIN analysis linked these programs through an inflammatory-vascular hub (BCL2, Cd9, CXCL12, Vegfa) and a glial node (AQP4) bridging myelination and tissue integrity (**Supplemental Fig. 8H**). Factor decomposition attributed Angiogenesis suppression to genotype (169 DEGs; **Supplemental Fig. 9B-C**), cytokine and microglial activation to injury (131 DEGs; **Supplemental Fig. 9D**) and suppression of Transmitter Response and Reuptake to the gene x injury interaction (52 DEGs; **Supplemental Fig. 9E**).

The cortical signature therefore integrates competing programs: neuroprotective and neurotrophic effectors (Bcl2 [35], Tgfb2 [36], Vegfa [37], Ppargc1a [38]) alongside chronic stress and secondary-injury markers (Ptgs2 [39], Xbp1 [40], Tnfrsf12a [41]), with reduced structural and synaptic gene expression (Aqp4 [42], Cntn1 [43], Nrxn1 [44]). To contextualize these findings, we examined *Hvcn1* and *Cybb* expression with age and injury and quantified brain immune and redox phenotypes at 18 months (**Supplemental Fig. 10**). *Hvcn1* rose with age in cortex and hippocampus (**Supplemental Fig. 10A-B**), with a parallel *Cybb* increase in hippocampus and a cortical trend (**Supplemental Fig. 10C-D**), identifying age as a major regulator of brain Hv1-NOX2 signaling. At the 18-month post-injury endpoint, CD45^int^CD11b^+^Ly6C^-^ microglia counts were significantly increased and peaked in Hv1KO/TBI mice (**Supplemental Fig. 10E**), whereas infiltrated myeloid cells and lymphocytes were unchanged (**Supplemental Fig. 10F-G**), indicating chronic microgliosis without peripheral infiltration.

Taken together, Hv1-deficient mice retained social and affective resilience relative to WT/TBI, without improvement in cognitive or locomotor functions. Cortical transcriptomic profiling revealed a response that combined neuroprotective signaling with concurrent chronic-stress and impaired structural-maintenance programs. Brain Hvcn1 expression increased with age, while Hv1KO/TBI cortex showed chronic microgliosis without peripheral infiltration. Hv1 deficiency thus yields a modest, mixed central phenotype that does not offset the systemic immune deterioration.

### Circulating plasma factors from chronic TBI and Hv1KO mice propagate systemic immune dysregulation and brain microgliosis in naïve recipients

To determine whether circulating factors contribute to the long-term propagation of systemic immune dysfunction after TBI, plasma collected from WT/Sham, WT/TBI, Hv1KO/Sham, or Hv1KO/TBI mice at 18 months post-injury was intravenously transferred into young adult WT recipients, followed by immunophenotypic and functional profiling (**Fig. 9A**). To provide cellular and functional validation of our transcriptomic datasets, flow cytometric readouts were selected based on genotype- and chronic TBI-dependent DEGs and pathway enrichment signatures identified through NanoString profiling, enabling cell-specific assessment of the most strongly altered innate immune pathways. Plasma transfer did not alter locomotor activity or anxiety-like behavior in the open field test (**Fig. 9B-C**), suggesting that short-term exposure to chronic TBI plasma was insufficient to produce overt behavioral abnormalities, thereby enabling interpretation of the observed immune phenotypes without confounding effects of neurological dysfunction.

**Fig. 9.**
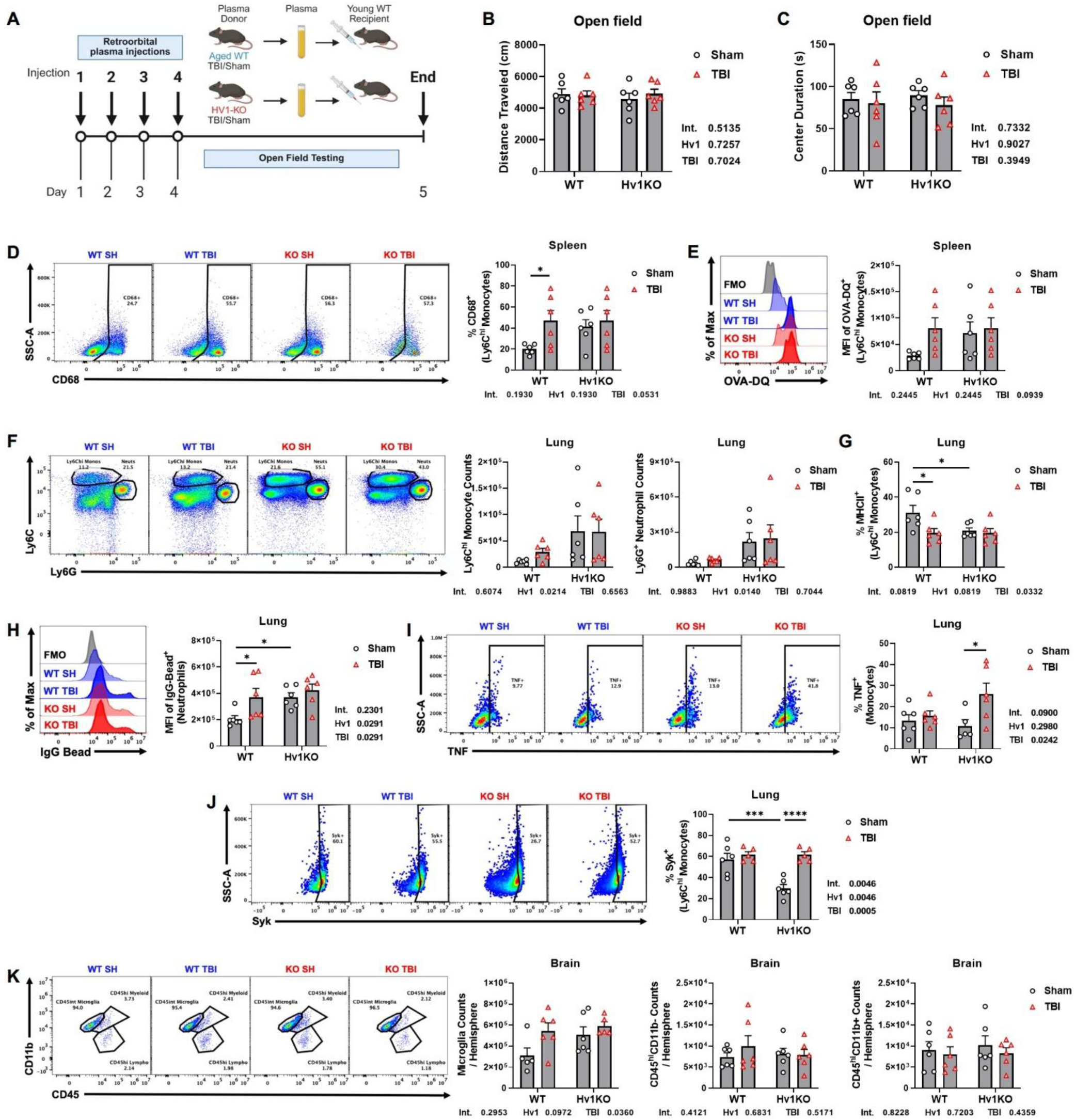
Plasma factors from chronic TBI and Hv1KO mice differentially shape immune responses in young adult WT recipients. **(A)** Experimental scheme of plasma transfer and post-injection immunophenotype and functional profiling. **(B-C)** Mice showed no difference of behaviors in open field test between groups. **(D-E)** Chronic TBI plasma derived from WT/TBI or Hv1KO mice promoted a more phagocytic and proteolytically active phenotype in splenic Ly6Chi monocytes, evidenced by increased CD68 expression and DQ-OVA processing relative to WT/Sham plasma. **(F)** Quantification of pulmonary Ly6C^hi^ monocyte and Ly6G+ neutrophil counts following injection of WT/Sham, WT/TBI, Hv1KO/Sham, or Hv1KO/TBI plasma into young adult WT mice, showing a significant main effect of Hv1 genotype on both monocyte and neutrophil populations. **(G)** In pulmonary Ly6C^hi^ monocytes of young adult mice, MHCII signaling was down-regulated by plasma from WT/TBI or Hv1KO. **(H)** Plasma injection from WT/TBI or Hv1KO (no difference between TBI and Sham) amplified IgG signaling in pulmonary neutrophils, indicating enhanced Fc-dependent uptake. **(I)** Chronic Hv1KO/TBI plasma upregulated TNF signal in monocytes of young adult mouse lungs. **(J)** Syk expression was exclusively reduced by Hv1KO/Sham plasma. **(K)** Increased microglia counts were detected in mouse brain after plasma injection of WT/TBI or Hv1KO (Sham or TBI) compared to WT/Sham, without alteration in infiltrated myeloid cells or lymphocytes. Sample sizes: N=5-6 per group. Statistical analyses: two-way ANOVA followed by Tukey’s post hoc test. *p≤0.05, **p≤0.01, ***p≤0.001, ****p≤0.0001.

In the spleen, plasma from both WT/TBI and Hv1KO donor mice promoted a more activated phagocyte phenotype, providing cellular-level validation of the immune pathways identified through transcriptomic analyses. Splenic Ly6C^hi^ monocytes exhibited increased CD68 protein expression and enhanced DQ-OVA proteolytic processing compared with recipients of WT/Sham plasma, indicating elevated phagocytic and lysosomal activity (**Fig. 9D-E**). In the lung, plasma transfer significantly altered innate immune populations and signaling pathways. Two-way ANOVA revealed a significant main effect of donor Hv1 genotype on pulmonary Ly6C^hi^ monocyte and Ly6G^+^ neutrophil abundance (**Fig. 9F**). Furthermore, MHCII expression in pulmonary Ly6C^hi^ monocytes was reduced following administration of WT/TBI or Hv1KO-derived plasma, suggesting suppression of antigen presentation pathways (**Fig. 9G**). In contrast, pulmonary neutrophils displayed enhanced IgG-associated signaling after exposure to WT/TBI or Hv1KO plasma regardless of injury status, consistent with increased Fc receptor-mediated uptake or immune complex engagement (**Fig. 9H**).

Additional evidence of differential immune programming was observed within the lung monocyte compartment. Plasma derived from Hv1KO/TBI mice selectively increased TNF signaling in pulmonary monocytes (**Fig. 9I**), whereas Syk expression was uniquely reduced following transfer of Hv1KO/Sham plasma (**Fig. 9J**). These findings indicate that chronic Hv1 deficiency and TBI generate distinct circulating immune signals capable of reprogramming peripheral myeloid cell function in recipient animals.

Within the CNS, plasma transfer was sufficient to alter resident microglial populations. Recipients of WT/TBI plasma or Hv1KO-derived plasma (Sham or TBI) exhibited increased microglial counts relative to mice receiving WT/Sham plasma (**Fig. 9K**). Notably, these changes occurred in the absence of detectable alterations in infiltrating myeloid cells or lymphocyte populations, suggesting that circulating factors primarily influenced resident microglial responses rather than peripheral leukocyte recruitment. Collectively, these findings demonstrate that chronic TBI and Hv1 deficiency generate circulating factors that remain biologically active months after injury and are sufficient to modulate immune phenotypes in the spleen, lung, and brain of otherwise healthy young adult recipients.

## Discussion

The present study, representing one of the longest longitudinal assessments of TBI (18 months post-injury), demonstrates that Hv1 deficiency preserves long-term neuroprotection but incurs a substantial systemic immune cost. The most striking finding was the progressive increase in mortality among Hv1KO/TBI mice during the chronic phase. Baseline Hv1 deficiency was also associated with weight loss and splenomegaly, pointing to broader systemic consequences. Evidence of spleen-restricted bacterial accumulation, impaired gut barrier and mucus integrity, and plasma transfer-induced immune alterations collectively implicates dysfunction of the gut-spleen axis as a potential driver of chronic immune dysregulation. Consistent with this, transcriptomic analyses revealed distinct organ-specific immune programs in the spleen, lung, and liver rather than a uniform systemic response, whereas cortical transcriptomic and behavioral analyses confirmed sustained neuroprotection. Finally, plasma transfer experiments demonstrated that chronic TBI- and Hv1 deficiency-associated circulating factors are sufficient to propagate immune alterations in peripheral organs and the brain, identifying blood-borne mediators as a potential mechanism underlying persistent systemic immune dysfunction.

A defining feature of the peripheral phenotype was that Hv1KO spleens were already abnormal at baseline. Even in the absence of injury, Hv1KO/Sham animals exhibited upregulation of pathogen-response, complement, and myeloid differentiation programs relative to WT/Sham controls, indicating a chronically activated immune state independent of TBI. Following chronic TBI, Hv1 deficiency produced a strikingly paradoxical transcriptional profile: upstream nucleic acid-sensing pathways were further activated yet type I interferon output was diminished; pro-inflammatory signaling was amplified, whereas antibacterial pathway activity was reduced. This disconnect between proximal immune sensing and downstream effector function resembles the immune landscape described in chronic inflammatory states such as post-sepsis immunosuppression [45] and persistent antigen exposure [46, 47], where signaling pathways remain engaged despite progressive impairment of antimicrobial defense. Although we avoid describing this phenotype as bona fide immune exhaustion in the absence of direct bactericidal or functional assays, its signaling architecture is consistent with an exhaustion-like state characterized by preserved immune activation but declining terminal effector capacity.

A plausible driver of this sustained immune burden is chronic translocation of gut-derived bacteria across a compromised intestinal barrier. Persistent low-level microbial exposure is well recognized as a stimulus for progressive immune dysfunction across both phagocyte and lymphocyte populations [47–49]. Likewise, chronic activation of NOX2-deficient phagocytes gives rise to the granulomatous, functionally hyporesponsive phenotype observed in chronic granulomatous disease (CGD) [50]. Although the present model differs from classical CGD in both mechanism and severity, this conceptual parallel supports the view that Hv1 deficiency promotes a gradual, antigen-driven decline in immune competence rather than an abrupt failure of host defense.

The extent of these peripheral immune abnormalities is also likely underestimated. Because the 18-month cohort consisted only of animals that survived long-term, it likely represents the more resilient end of the disease spectrum rather than the full range of pathological severity. Animals that died earlier may have developed more severe splenic dysfunction, greater compromise of intestinal barrier integrity, and increased systemic bacterial translocation. Consequently, survivor bias is likely obscured by the true magnitude of the peripheral immune deficits associated with Hv1 deficiency following chronic TBI.

The molecular changes observed in the chronically injured Hv1-deficient gut provide a plausible explanation for the spleen-restricted bacterial accumulation. Following disruption of the intestinal barrier, translocated bacteria enter the portal circulation, where they are first filtered by Kupffer cells in the liver before reaching the spleen for terminal clearance by red-pulp and marginal-zone phagocytes [51, 52]. Experimental TBI disrupts intestinal barrier integrity, alters the gut microbiota, and promotes bacterial translocation, with effects that persist well beyond the acute phase [53–56]. In WT animals, splenic phagocytes eliminate these bacteria through NOX2-dependent oxidative burst, whereas Hv1 deficiency is expected to impair this antimicrobial function. The absence of detectable bacterial accumulation in the liver suggests that hepatic clearance remains largely intact, while the splenic bacterial burden is consistent with impaired downstream clearance. Future studies incorporating gut microbiome profiling and bacterial translocation tracing will be needed to directly test this gut–liver–spleen axis.

Additional evidence for persistent systemic dysregulation comes from the plasma transfer experiments. Plasma from chronic TBI or Hv1-deficient donors was sufficient to reproduce key peripheral phenotypes in naïve young adult WT recipients, including enhanced splenic monocyte phagocytic and proteolytic activity, altered pulmonary monocyte and neutrophil signaling, and increased brain microgliosis without detectable immune-cell infiltration. These findings indicate that chronic circulating factors are sufficient to reprogram myeloid function across multiple organs. The identity of these factors remains unknown but could include translocated microbial products [3], inflammatory cytokines [57], host-derived damage-associated molecules such as mitochondrial DNA and formyl peptides, or microbial cell-free DNA [58, 59]. Because donor plasma was dialyzed before transfer, the transferable activity is likely carried by components larger than 3.5 kDa, such as proteins, peptides, nucleic acid-containing complexes, extracellular vesicles, or microbial products, rather than small soluble metabolites..

The three peripheral organs profiled here showed distinct transcriptional signatures, reflecting their characteristic immune cell composition, baseline functional programming, regenerative capacity, and aging trajectory. All the factors together, reshape each organ’s homeostatic balance according to its physiological role under a single Hvcn1 channel deletion. The spleen relies on specialized resident macrophages, including marginal-zone, marginal-metallophilic, and red-pulp macrophages, to eliminate blood-borne pathogens and rapidly initiate adaptive immunity [51, 52, 60]. These cells depend on NOX2-mediated oxidative burst for bacterial killing, but their marginal-zone architecture is highly susceptible to chronic infection, inflammation, and aging, leading to impaired pathogen clearance with limited structural recovery [61–65]. As the terminal filter for encapsulated bacteria that escape hepatic clearance, splenic dysfunction ultimately results in bacterial accumulation, consistent with our findings [66]. In contrast, the lung is populated by tissue-resident alveolar macrophages maintained in a restrained inflammatory state, together with interstitial macrophages and recruited neutrophils, to preserve alveolar integrity during constant environmental exposure [67, 68]. Aging shifts this balance toward inflammatory monocyte-derived macrophages and dysregulated alveolar macrophage responses [69, 70]. Accordingly, the inflammatory signature observed in Hv1KO/TBI lungs, including increased chemokine, complement, and TLR pathway activity with elevated CD11b⁺ infiltration, likely reflects chronic systemic inflammation rather than intrinsic pulmonary clearance failure. The liver represents a distinct immune-metabolic compartment. As the first organ exposed to portal venous blood, it combines Kupffer cell phagocytosis with hepatocyte acute-phase responses, sinusoidal endothelial scavenging, immune tolerance, and remarkable regenerative capacity [71–73]. Although we observed metabolic suppression and a pro-inflammatory transcriptional shift, the absence of cGAS-STING-IFN activation or bacterial accumulation indicates preserved Kupffer-cell clearance. We propose that this resilience reflects continuous Kupffer-cell replenishment through local proliferation and monocyte recruitment, together with hepatocyte renewal and sinusoidal remodeling [72, 73]. By comparison, splenic red-pulp macrophages have limited self-renewal capacity that further declines with aging [74]. This difference may explain why the liver maintains antibacterial function during chronic disease whereas the spleen progresses to clearance failure, suggesting that enhancing splenic phagocyte renewal could mitigate the gut–spleen axis driving late mortality.

Consistent with these architectural differences, cell-type deconvolution and flow cytometry revealed distinct organ-specific leukocyte responses that help explain why only the spleen progressed to antibacterial failure. The spleen of Hv1KO/TBI mice showed accumulation of CD11b⁺ myeloid cells with increased neutrophil and macrophage signatures but reduced B-cell and overall CD45 enrichment, indicating chronic myeloid activation alongside weakening adaptive immunity. In contrast, the lung maintained expanded myeloid and dendritic populations with relatively preserved cytotoxic T-cell enrichment, supporting effective bacterial control despite heightened inflammation. The liver exhibited increased CD45⁺ cells, macrophages, and neutrophils only when Hv1 deficiency and TBI coincided, consistent with an inducible, regenerable myeloid barrier. Together, these findings suggest that the spleen’s fixed architecture and limited regenerative capacity, rather than inflammatory intensity alone, make it the weakest link in systemic antibacterial defense during chronic Hv1 deficiency.

Social novelty preference, novelty-suppressed feeding latency, and other cognitive measures were preserved or improved in Hv1KO/TBI mice, consistent with the established neuroprotective effects of Hv1 deficiency from the acute through chronic phases in young adults [10, 11, 14, 16]. However, these behavioral benefits were not fully reflected at the molecular level. Cortical microgliosis persisted in Hv1KO/TBI mice without increased peripheral immune cell infiltration, suggesting expansion of resident microglia rather than recruitment of circulating cells. At 18 months, NOX2 protein and intracellular ROS levels were unchanged, indicating that chronic oxidative stress may arise from alternative ROS sources or reduced NOX2 dependence. Consistent with this, transcriptomic analysis showed persistent enrichment of oxidative stress and tissue integrity pathways. The age-related increase in Hvcn1 and Cybb expression in the wild-type cortex and hippocampus further suggests that Hv1-NOX2 signaling becomes more prominent with aging, highlighting that long-term Hv1 deficiency cannot be fully understood by young adults alone [4, 75, 76]. Together, these findings extend previous work by supporting long-term, microglia-targeted Hv1 inhibition as a potential therapeutic strategy for TBI, particularly in older individuals at greater risk of neurodegeneration.

Future studies should use cell-specific Hv1 knockout models and include both sexes to clarify cellular and sex-dependent mechanisms. Molecular profiling of plasma, gut microbiota analyses, metabolomics, and functional studies of splenic phagocytes will help identify circulating mediators and further define the gut-spleen axis and immune exhaustion. Therapeutically, cell-and time-specific Hv1 modulation may offer greater benefit than systemic inhibition. Plasma 16S rRNA and IFNβ are promising biomarkers, while gut-targeted therapies may provide complementary benefit. Defining the optimal therapeutic window will be critical for clinical translation.

## Conclusions

In summary, Hv1 emerges as a context-dependent immune regulator, whose deficiency produces adverse consequences of compromised systemic immune integrity and reduced survival despite preserved neuroprotection after long-term TBI. Transcriptomic profiling reveals aberrant splenic immune responses along with damaged gut barriers driven by the interaction of Hv1KO and TBI, providing potential insights for the underlying mechanisms. The hypothesis is supported by the circulating immune-modulating activity in naïve young WT mice after plasma transfer. These findings argue for cell-restricted, time-specified Hv1 modulation strategies and identify the gut-spleen axis as a critical target for adjunctive intervention in chronic TBI.

## Conflict of interest statement

The authors have no conflicts of interest to declare.

## Authors’ contributions

Z.L. performed the behavioral tests, RT-qPCR experiments, NanoString analysis, and wrote the manuscript. R.K. conducted in vivo plasma injection and flow cytometry experiments. Y.L. and K.B. contributed to sample collection and RNA extraction. C.R.S. and P.J.D. assisted with the plasma transfer studies. J.H. performed CCI surgeries. R.M.R. conceived this study, conducted flow cytometry experiments and plasma transfer studies, supervised the project, and revised the manuscript. J.W. contributed to studying conception and design and revised the manuscript.

## Acknowledgments

We thank H. Li for assistance with animal care and blood preparation.

## Data availability

The datasets used and/or analyzed during the current study are available from the corresponding author on reasonable request.

## Declarations

### Funding

This work was supported by grants from the NIH R01AG077541 (J.W.), R01NS145443 (J.W.), R01NS110825 (J.W.), and NIH R00NS116032 (to R.M.R.).

### Ethics approval and consent to participate

Animal experiments were carried out according to experimental protocols approved by the Animal Care and Use Committee (IACUC) of the University of Maryland School of Medicine and University of Texas Health Science Center at Houston Institutional.

### Consent for publication

Not applicable.

## Notes

### Competing Interest Statement

The authors have declared no competing interest.

