## Supplemental Figures for "Hv1 proton channel is essential for splenic antibacterial defense and immune homeostasis during the chronic phase of traumatic brain injury in male mice"

**Supplementary Information**

Supplemental Information includes ten Supplemental figures and figure legends.

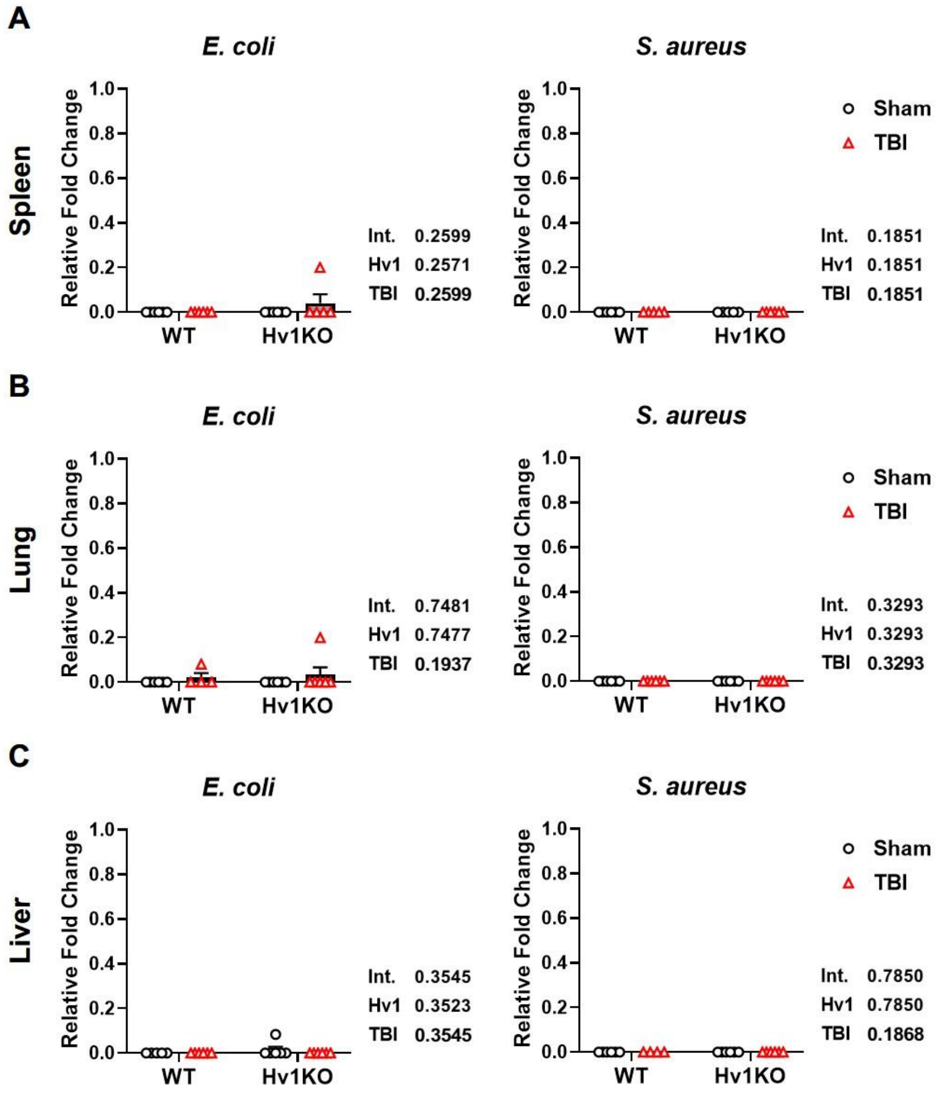


**Supplemental Fig. 1. *E. coli* and *S. aureus* signals remain near the detection floor in spleen, lung, and liver.** (**A**) In spleen, *E. coli* (A) and *S. aureus* (B) signals were near or below detection with no group differences. (**B-C**) Comparable results in lung (B) and liver (C). Sample sizes: n=5–6 mice/group. Statistical analyses: Two-way ANOVA with Šídák-corrected simple-effects comparisons (A–C) with p-values representing interaction (Int.), *Hvcn1* knock-out (Hv1), and TBI main-effect shown beside each panel.

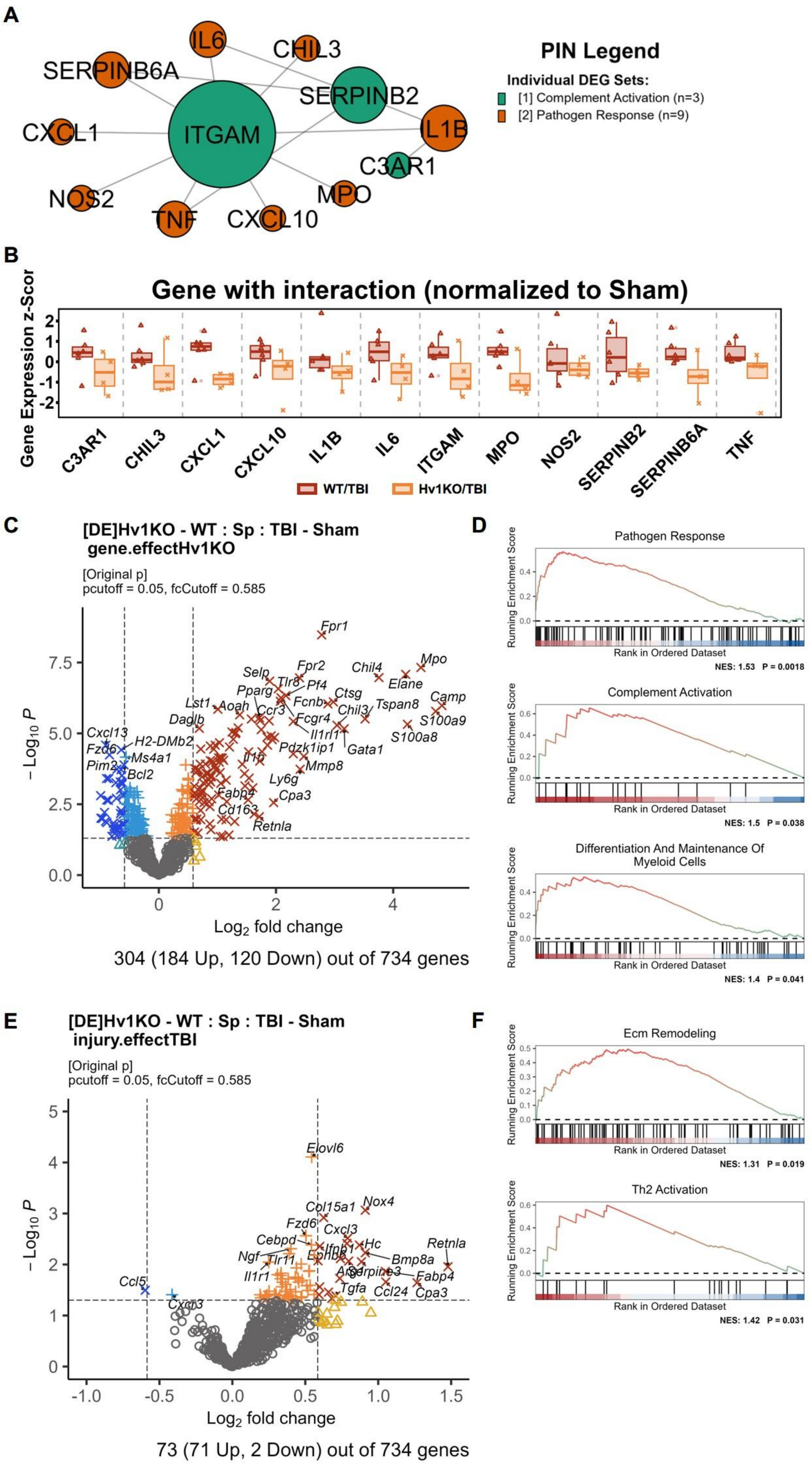
A
PIN Legend
Individual DEG Sets:
[1] Complement Activation (n=3)
[2] Pathogen Response (n=9)
B
Gene with interaction (normalized to Sham)
Gene Expression z-Scor
WT/TBI
Hv1KO/TBI
C
[DE]Hv1KO - WT : Sp : TBI - Sham
gene.effectHv1KO
[Original p]
pcutoff = 0.05, fcCutoff = 0.585
Log2 fold change
304 (184 Up, 120 Down) out of 734 genes
D
Pathogen Response
NES: 1.53 P = 0.0018
Complement Activation
NES: 1.5 P = 0.038
Differentiation And Maintenance Of Myeloid Cells
NES: 1.4 P = 0.041
Running Enrichment Score
Rank in Ordered Dataset
E
[DE]Hv1KO - WT : Sp : TBI - Sham
injury.effectTBI
[Original p]
pcutoff = 0.05, fcCutoff = 0.585
Log2 fold change
73 (71 Up, 2 Down) out of 734 genes
F
Ecm Remodeling
NES: 1.31 P = 0.019
Th2 Activation
NES: 1.42 P = 0.031

**Supplemental Fig. 2. Two-factor (genotype × injury) differential expression and pathway analysis in the spleen. (A)** Protein-protein interaction network (PIN) of genes with a significant Hv1KO × TBI interaction, restricted to the "Complement Activation" (n = 3, green) and "Pathogen Response" (n = 9, orange) modules. Node color denotes DEG-set membership and node size reflects connectivity within the network; *ITGAM* and *SERPINB2* are the principal hubs. **(B)** Expression z-scores (normalized to Sham) for the same interaction genes; the WT/TBI group (dark-red) showed higher gene expression than the Hv1KO/TBI group (orange) across most genes. **(C-D)** Genotype main effect (Hv1KO vs WT) produced the volcano plot (C) demonstrating 304 significant DEGs out of 734 genes, and GSEA (D) showing positive enrichment of "Pathogen Response", "Complement Activation“, and "Differentiation and Maintenance of Myeloid Cells". **(E-F)** Injury main effect (TBI vs Sham) generated the volcano plot (E): 73/734 DEGs (71 up, 2 down); and GSEA (F): positive enrichment of "Ecm Remodeling" and "Th2 Activation". DEGs were significant at $p<0.05$. Sample sizes: WT/Sham, n=6; Hv1KO/Sham, n=4; WT/TBI, n=6; Hv1KO/TBI, n=4.

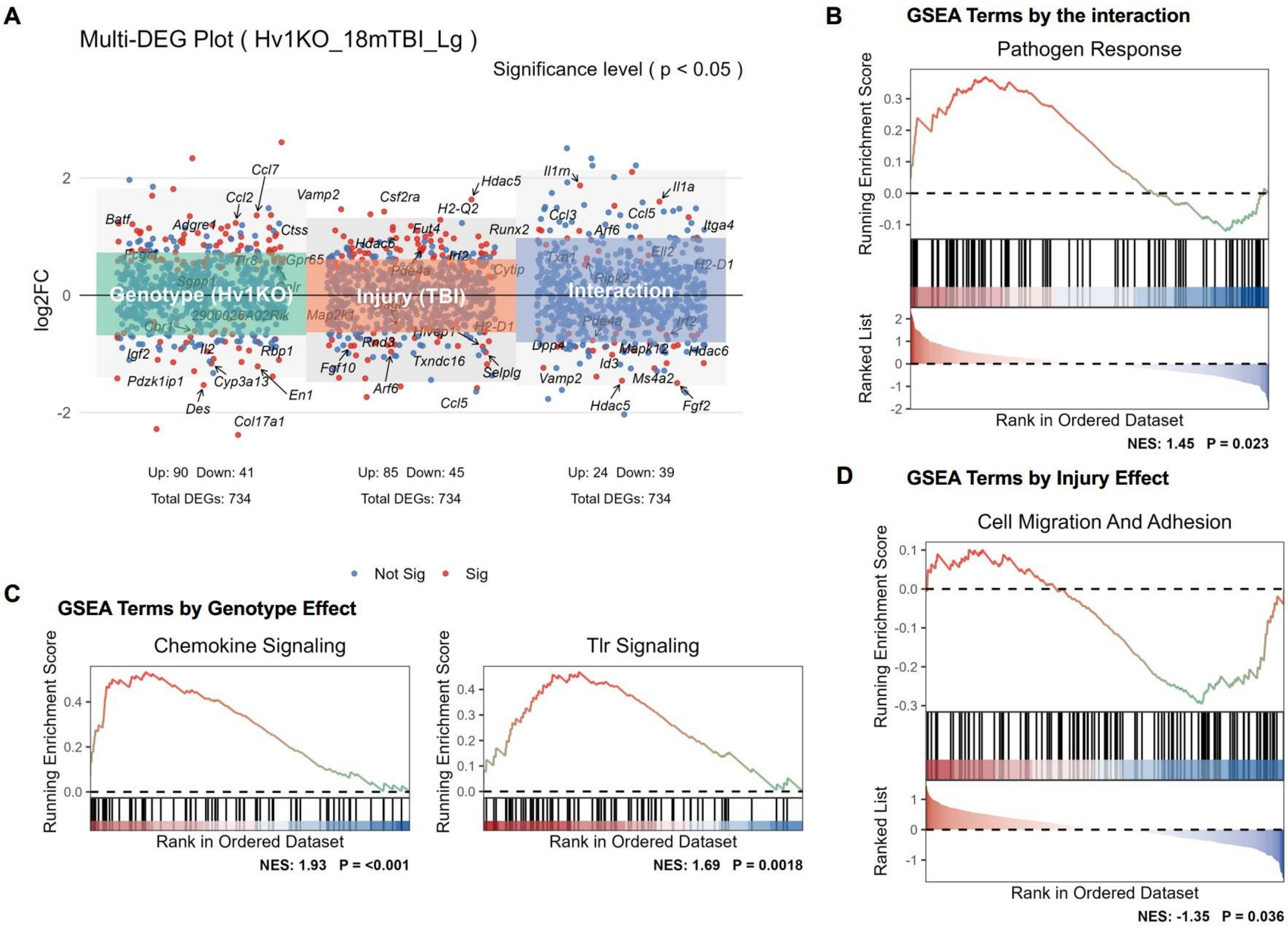


**Supplemental Fig. 3. Two-factor (genotype × injury) differential expression and pathway analysis in the lung. (A)** Multi-DEG plot of per-gene $\log_2$ fold change for each contrast (genotype, Hv1KO vs WT; injury, TBI vs Sham; interaction); red points are significant ($p < 0.05$), blue are non-significant. **(B)** GSEA of the interaction term shows positive enrichment of "Pathogen Response". **(C)** Genotype effect positively enriches "Chemokine Signaling" and "Tlr Signaling". **(D)** Injury effect negatively enriches "Cell Migration and Adhesion". Group sizes: WT/Sham n = 4; Hv1KO/Sham n = 6; WT/TBI n = 5; Hv1KO/TBI n = 6.

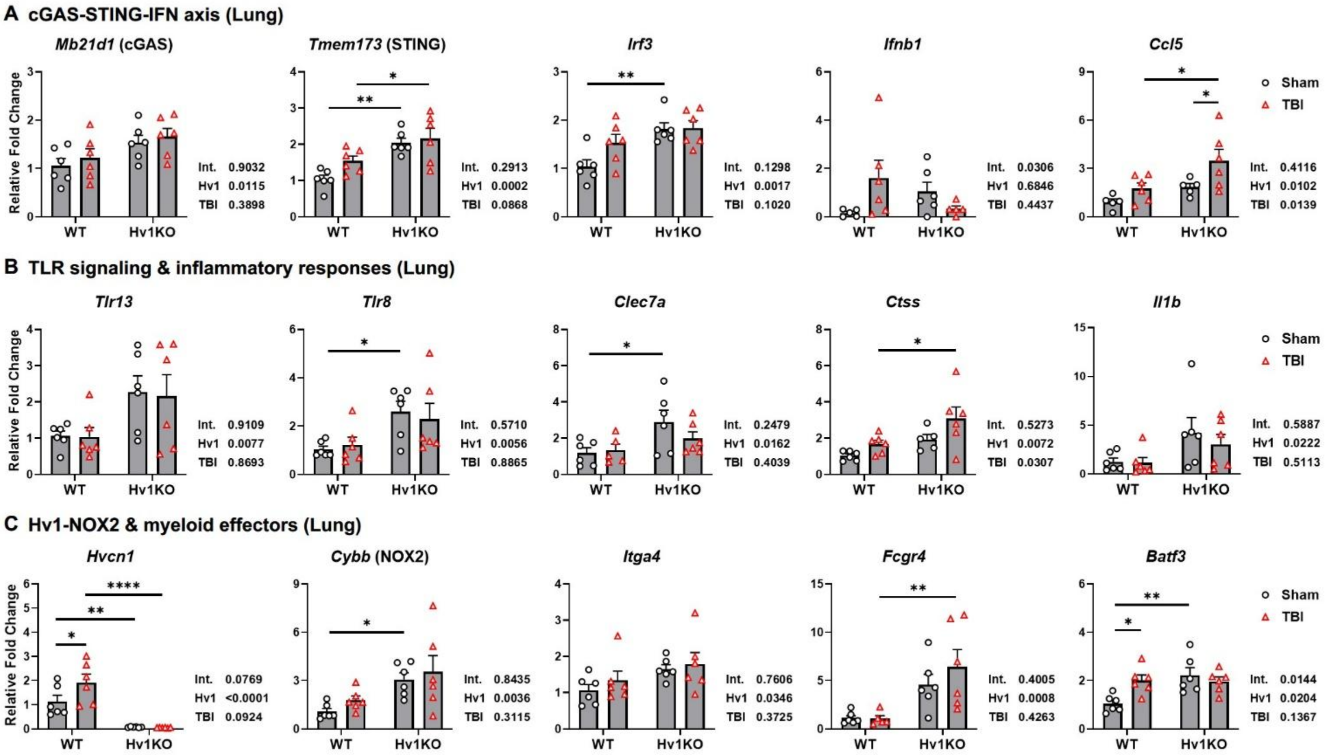


**Supplemental Fig. 4. Hv1 deficiency reshapes innate-immune gene expression in the lung and alters the transcriptional response to chronic TBI.** Relative mRNA (RT-qPCR) in lungs of WT and Hv1KO mice at 18 months after Sham or TBI, grouped into three functional modules. **(A)** Cytosolic DNA sensing and type I IFN: *Mb21d1* (cGAS), *Tmem173* (STING), *Irf3*, *Ifnb1*, and the IFN-responsive chemokine *Ccl5*. **(B)** TLR/CLR sensing and inflammatory output: *Tlr13*, *Tlr8*, *Clec7a* (Dectin-1), *Ctss* (cathepsin S), and *Il1b*. **(C)** Hv1-NOX2 redox and myeloid effectors: *Hvcn1* (Hv1), *Cybb* (NOX2), *Itga4*, *Fcgr4* (FcγRIV), and *Batf3*. Sample sizes: n=5-6 mice/group. Statistical analyses: Two-way ANOVA with Šídák-corrected simple-effects comparisons (A-C) with p-values representing interaction (Int.), *Hvcn1* knock-out (Hv1), and TBI main-effect shown beside each panel. *p≤0.05, **p≤0.01, ***p≤0.001, ****p≤0.0001.

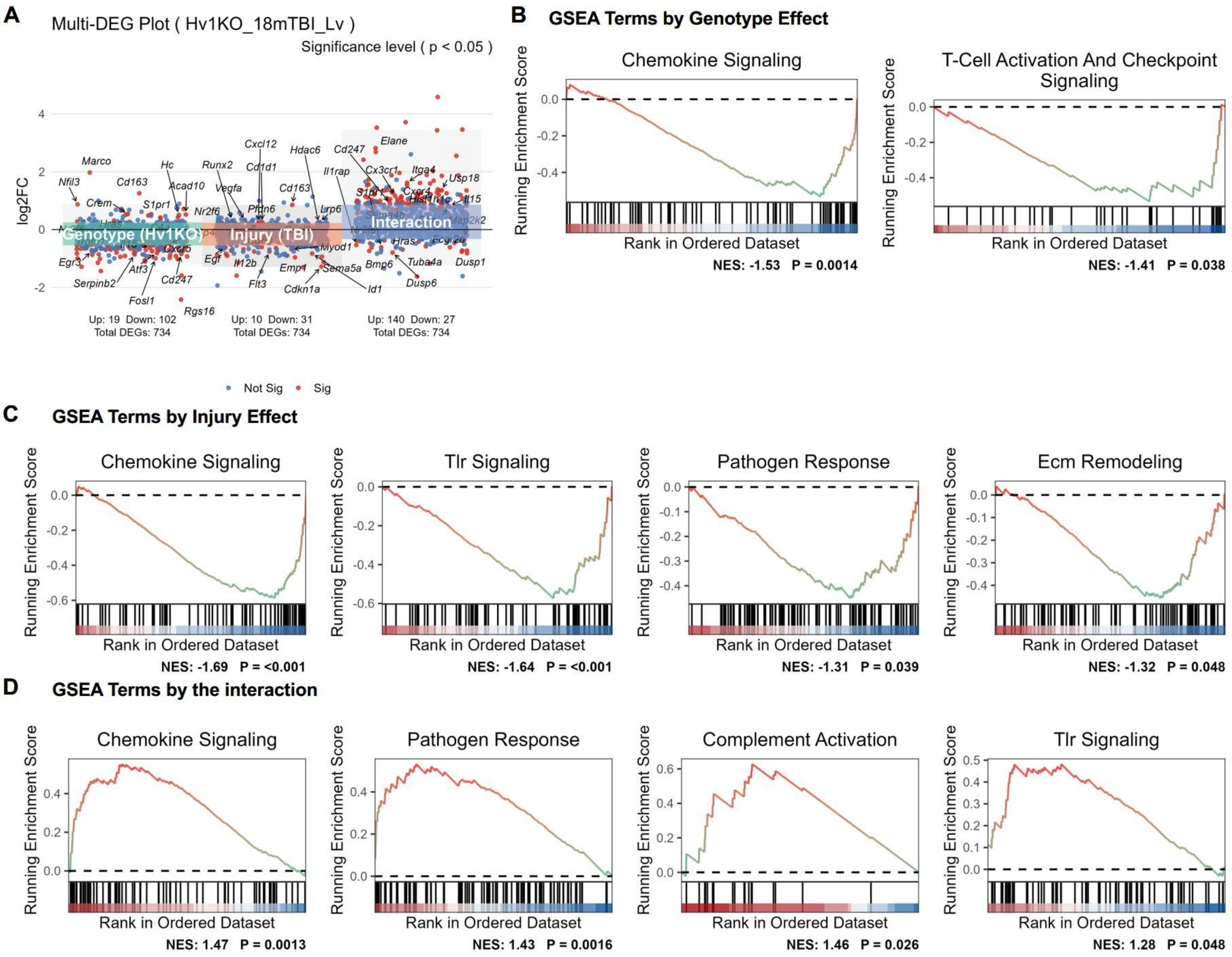


**Supplemental Fig. 5. The genotype × injury interaction dominates the hepatic transcriptional response, activating chemokine, complement, and innate-immune programs in the liver. (A)** Multi-DEG plot of per-gene $\log_2$ fold change for each contrast (genotype, Hv1KO vs WT; injury, TBI vs Sham; interaction); red points are significant ($p < 0.05$), blue non-significant. Panel-based GSEA for each contrast: **(B)** Genotype effect suppresses "Chemokine Signaling" and "T-Cell Activation and Checkpoint Signaling". **(C)** Injury effect suppresses "Chemokine Signaling", "Tlr Signaling", "Pathogen Response", and "Ecm Remodeling". **(D)** Interaction activates "Chemokine Signaling", "Pathogen Response", "Complement Activation", and "Tlr Signaling". Sample sizes: WT/Sham, n=6; Hv1KO/Sham, n=6; WT/TBI, n=6; Hv1KO/TBI, n=6.

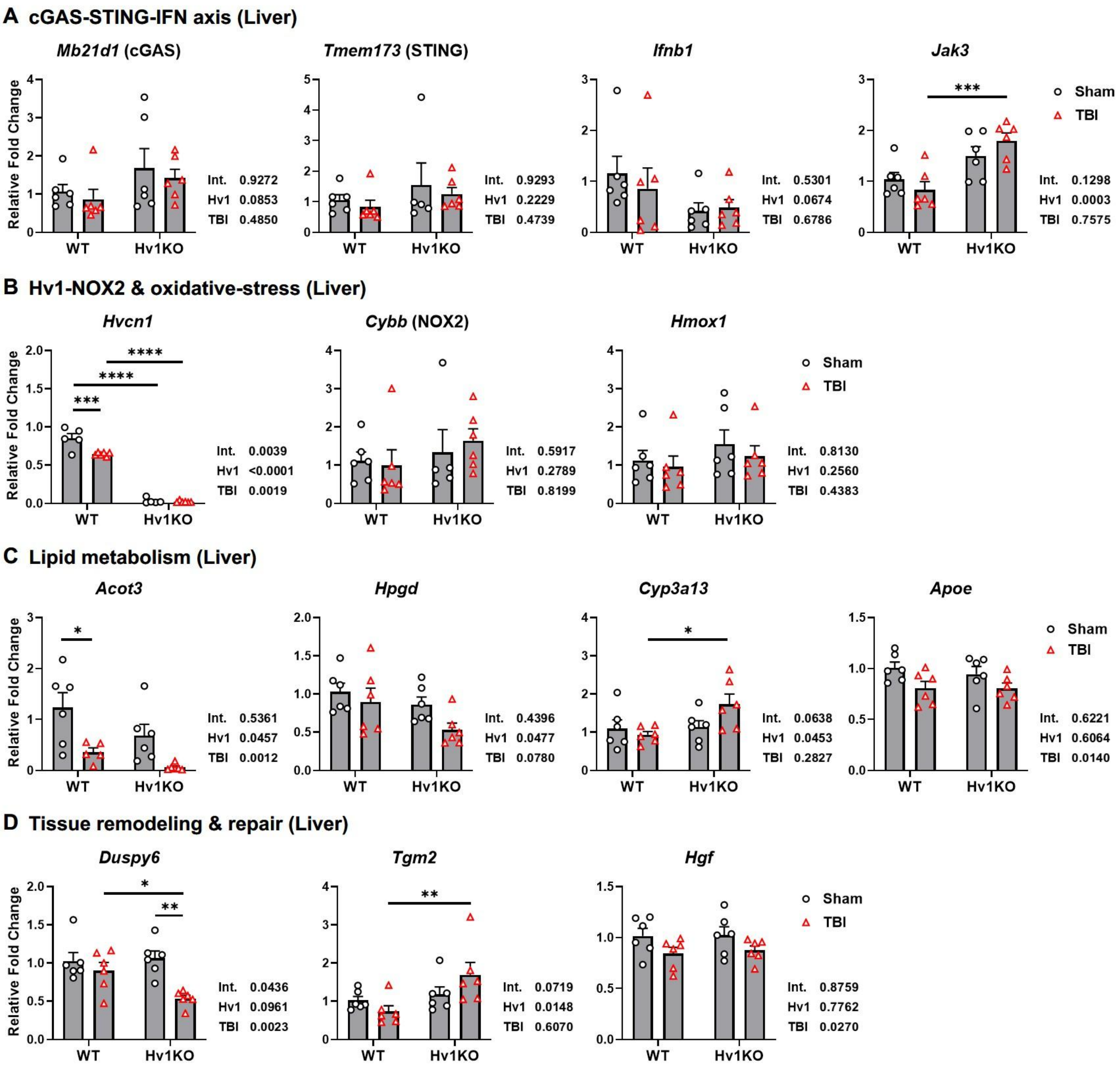
A cGAS-STING-IFN axis (Liver)
Mb21d1 (cGAS)
Int. 0.9272
Hv1 0.0853
TBI 0.4850
Tmem173 (STING)
Int. 0.9293
Hv1 0.2229
TBI 0.4739
Ifnb1
Int. 0.5301
Hv1 0.0674
TBI 0.6786
Jak3
***
Int. 0.1298
Hv1 0.0003
TBI 0.7575
Sham
TBI
Relative Fold Change
WT
Hv1KO
B Hv1-NOX2 & oxidative-stress (Liver)
Hvcn1
***
****
****
Int. 0.0039
Hv1 <0.0001
TBI 0.0019
Cybb (NOX2)
Int. 0.5917
Hv1 0.2789
TBI 0.8199
Hmox1
Int. 0.8130
Hv1 0.2560
TBI 0.4383
Sham
TBI
C Lipid metabolism (Liver)
Acot3
*
Int. 0.5361
Hv1 0.0457
TBI 0.0012
Hpgd
Int. 0.4396
Hv1 0.0477
TBI 0.0780
Cyp3a13
*
Int. 0.0638
Hv1 0.0453
TBI 0.2827
Apoe
Int. 0.6221
Hv1 0.6064
TBI 0.0140
Sham
TBI
D Tissue remodeling & repair (Liver)
Duspy6
*
**
Int. 0.0436
Hv1 0.0961
TBI 0.0023
Tgm2
**
Int. 0.0719
Hv1 0.0148
TBI 0.6070
Hgf
Int. 0.8759
Hv1 0.7762
TBI 0.0270
Sham
TBI

**Supplemental Fig. 6. Hv1 deficiency alters hepatic gene expression and partially reshapes the transcriptional response to chronic TBI.** Relative mRNA (RT-qPCR) in livers of WT and Hv1KO mice at 18 months after Sham or TBI, grouped into four functional modules. **(A)** cGAS-STING-IFN axis: *Mb21d1* (cGAS), *Tmem173* (STING), *Ifnb1*, and the cytokine-signaling kinase *Jak3*. **(B)** Hv1-NOX2 and oxidative stress: *Hvcn1* (Hv1), *Cybb* (NOX2), *Hmox1*. **(C)** Lipid and xenobiotic metabolism: *Acot3*, *Hpgd*, *Cyp3a13*, *Apoe*. **(D)** Tissue remodeling and repair: *Dusp6*, *Tgm2*, *Hgf*.. For each panel, Sample sizes: n=5-6 mice/group. Sample sizes: n=5–6 mice/group. Statistical analyses: Two-way ANOVA with Šídák-corrected simple-effects comparisons (A-D) with p-values representing interaction (Int.), *Hvcn1* knock-out (Hv1), and TBI main-effect shown beside each panel. *p≤0.05, **p≤0.01, ***p≤0.001, ****p≤0.0001.

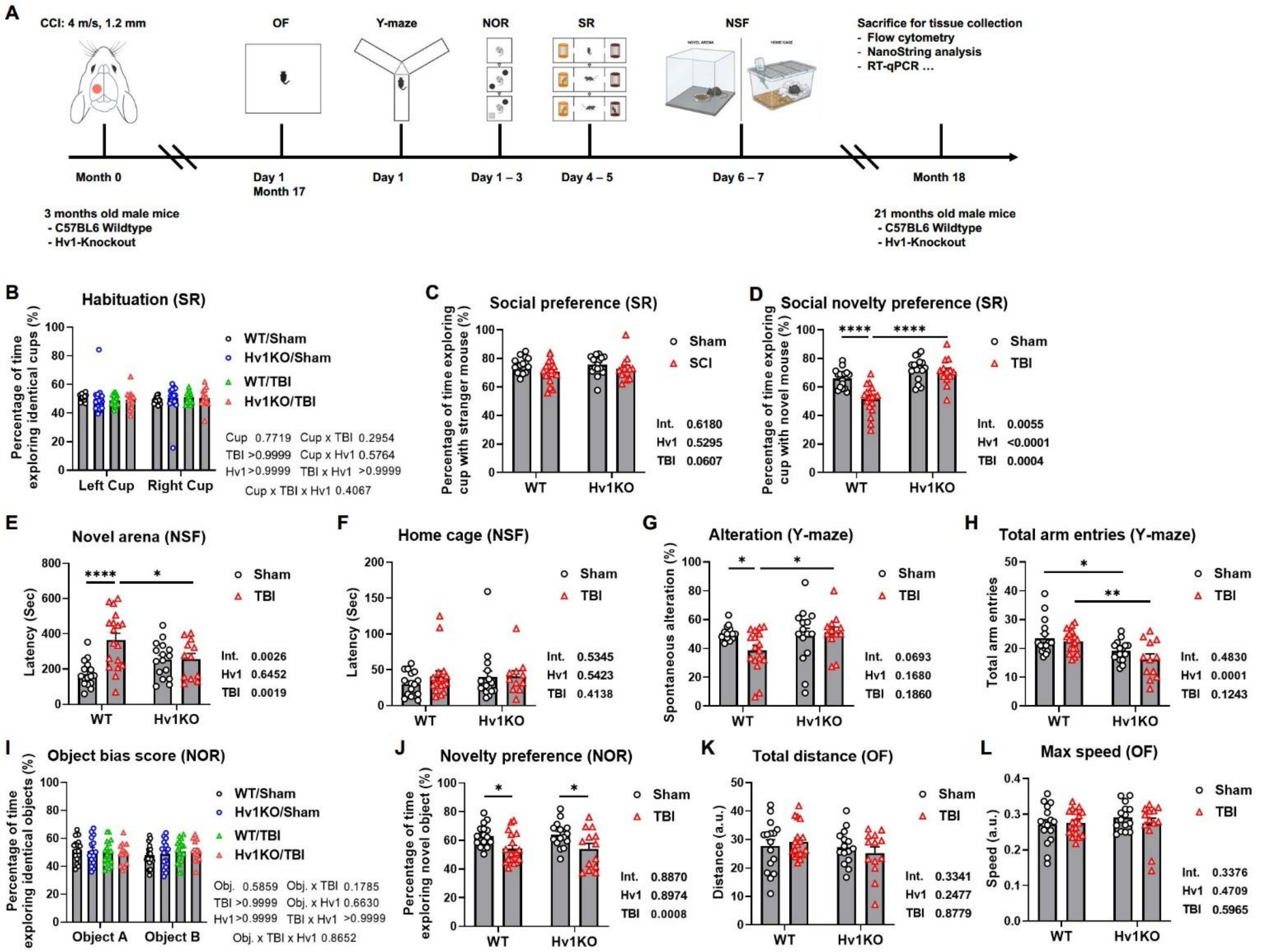
A
CCI: 4 m/s, 1.2 mm
OF
Y-maze
NOR
SR
NSF
Sacrifice for tissue collection
- Flow cytometry
- NanoString analysis
- RT-qPCR …
Month 0
Day 1
Month 17
Day 1
Day 1 – 3
Day 4 – 5
Day 6 – 7
Month 18
3 months old male mice
- C57BL6 Wildtype
- Hv1-Knockout
21 months old male mice
- C57BL6 Wildtype
- Hv1-Knockout
B Habituation (SR)
Percentage of time exploring identical cups (%)
WT/Sham
Hv1KO/Sham
WT/TBI
Hv1KO/TBI
Cup 0.7719 Cup x TBI 0.2954
TBI >0.9999 Cup x Hv1 0.5764
Hv1 >0.9999 TBI x Hv1 >0.9999
Cup x TBI x Hv1 0.4067
Left Cup
Right Cup
C Social preference (SR)
Percentage of time exploring cup with stranger mouse (%)
Sham
SCI
Int. 0.6180
Hv1 0.5295
TBI 0.0607
WT
Hv1KO
D Social novelty preference (SR)
Percentage of time exploring cup with novel mouse (%)
Sham
TBI
Int. 0.0055
Hv1 <0.0001
TBI 0.0004
E Novel arena (NSF)
Latency (Sec)
Int. 0.0026
Hv1 0.6452
TBI 0.0019
F Home cage (NSF)
Latency (Sec)
Int. 0.5345
Hv1 0.5423
TBI 0.4138
G Alteration (Y-maze)
Spontaneous alteration (%)
Int. 0.0693
Hv1 0.1680
TBI 0.1860
H Total arm entries (Y-maze)
Total arm entries
Int. 0.4830
Hv1 0.0001
TBI 0.1243
I Object bias score (NOR)
Percentage of time exploring identical objects (%)
Obj. 0.5859 Obj. x TBI 0.1785
TBI >0.9999 Obj. x Hv1 0.6630
Hv1 >0.9999 TBI x Hv1 >0.9999
Obj. x TBI x Hv1 0.8652
Object A
Object B
J Novelty preference (NOR)
Percentage of time exploring novel object (%)
Int. 0.8870
Hv1 0.8974
TBI 0.0008
K Total distance (OF)
Distance (a.u.)
Int. 0.3341
Hv1 0.2477
TBI 0.8779
L Max speed (OF)
Speed (a.u.)
Int. 0.3376
Hv1 0.4709
TBI 0.5965

**Supplemental Fig. 7. Hv1 deficiency confers partial behavioral resilience after chronic TBI. (A)** Experimental design: 3-month-old male WT (C57BL/6) and Hv1KO mice underwent CCI (4 m/s, 1.2 mm) or Sham, a behavioral battery at 17 months, and sacrifice at 18 months for flow cytometry, NanoString, and RT-qPCR. **(B-D)** Three-chamber social recognition (SR): habituation with no cup preference (B), social preference for the stranger-containing chamber (C), and social novelty preference (D), where Hv1KO/TBI mice retained preference for the novel mouse while WT/TBI did not. **(E-F)** Novelty-suppressed feeding (NSF): latency to feed in the home cage (E) and novel arena (F). **(G-H)** Y-maze: spontaneous alternation (G) and total arm entries (H). **(I-J)** Novel object recognition (NOR): object bias at baseline (I) and novelty preference (J). **(K-L)** Open field: distance traveled (K) and maximum speed (L). Sample sizes: WT/Sham, n=16; Hv1KO/Sham, n=16; WT/TBI, n=18; Hv1KO/TBI, n=13. Statistical analyses: three-way ANOVA followed by Šídák's multiple comparisons test for (C) and (J); Two-way ANOVA with Šídák-corrected simple-effects comparisons (C-H, J-L) with p-values representing interaction (Int.), *Hvcn1* knock-out (Hv1), and TBI main-effect shown beside each panel. *p<0.05, **p<0.01, ***p<0.001, ****p<0.0001.

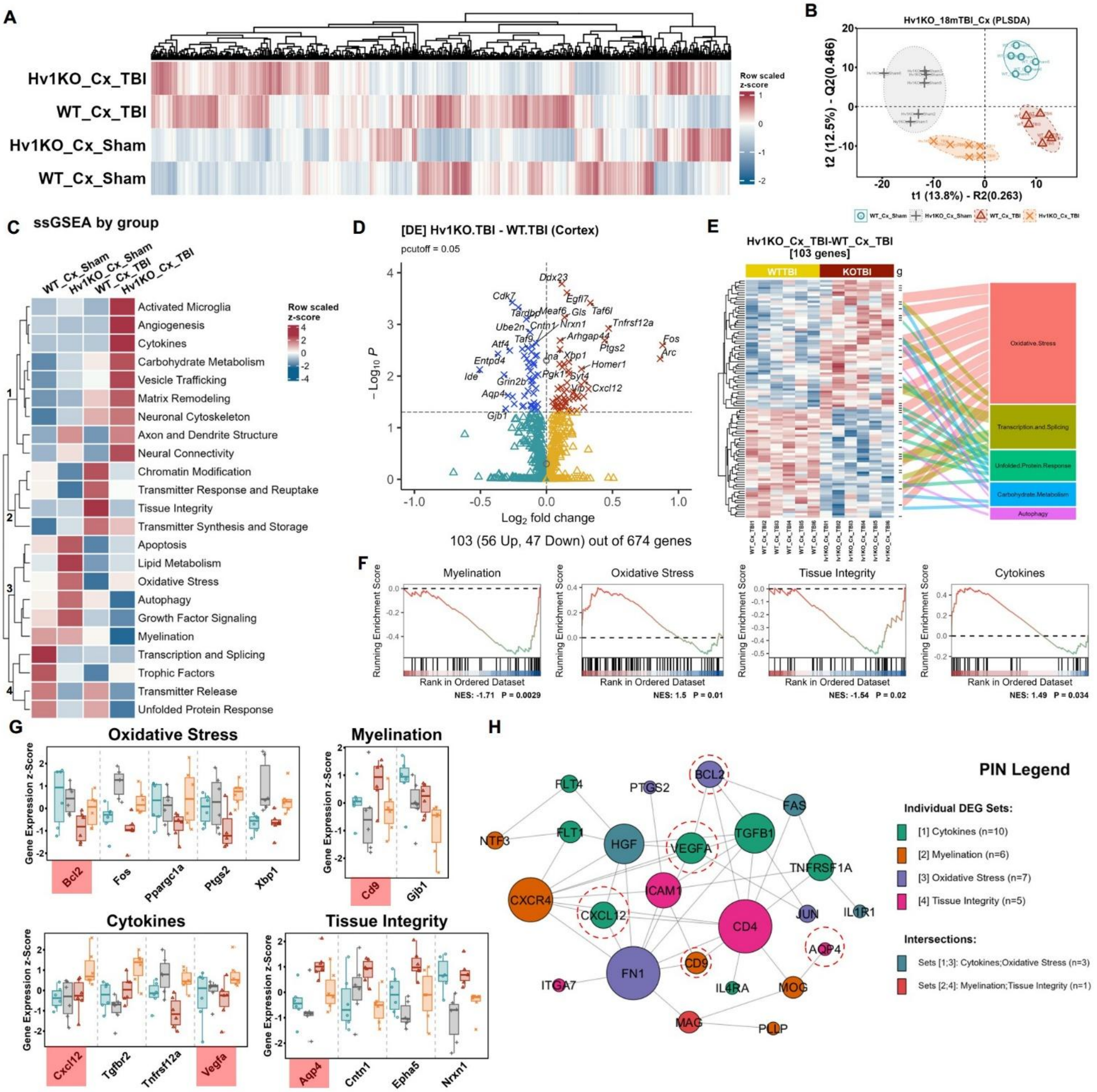
A
Hv1KO_Cx_TBI
WT_Cx_TBI
Hv1KO_Cx_Sham
WT_Cx_Sham
Row scaled z-score
B
Hv1KO_18mTBI_Cx (PLSDA)
t1 (13.8%) - R2(0.263)
t2 (12.5%) - Q2(0.466)
WT_Cx_Sham
Hv1KO_Cx_Sham
WT_Cx_TBI
Hv1KO_Cx_TBI
C
ssGSEA by group
WT_Cx_Sham
Hv1KO_Cx_Sham
WT_Cx_TBI
Hv1KO_Cx_TBI
Activated Microglia
Angiogenesis
Cytokines
Carbohydrate Metabolism
Vesicle Trafficking
Matrix Remodeling
Neuronal Cytoskeleton
Axon and Dendrite Structure
Neural Connectivity
Chromatin Modification
Transmitter Response and Reuptake
Tissue Integrity
Transmitter Synthesis and Storage
Apoptosis
Lipid Metabolism
Oxidative Stress
Autophagy
Growth Factor Signaling
Myelination
Transcription and Splicing
Trophic Factors
Transmitter Release
Unfolded Protein Response
D
[DE] Hv1KO.TBI - WT.TBI (Cortex)
pcutoff = 0.05
Log2 fold change
103 (56 Up, 47 Down) out of 674 genes
E
Hv1KO_Cx_TBI-WT_Cx_TBI
[103 genes]
WTTBI
KOTBI
Oxidative.Stress
Transcription.and.Splicing
Unfolded.Protein.Response
Carbohydrate.Metabolism
Autophagy
F
Myelination
Oxidative Stress
Tissue Integrity
Cytokines
Running Enrichment Score
Rank in Ordered Dataset
NES: -1.71 P = 0.0029
NES: 1.5 P = 0.01
NES: -1.54 P = 0.02
NES: 1.49 P = 0.034
G
Oxidative Stress
Bcl2
Fos
Ppargc1a
Ptgs2
Xbp1
Myelination
Cd9
Gjb1
Cytokines
Cxcl12
Tgfbr2
Tnfrsf12a
Vegfa
Tissue Integrity
Aqp4
Cntn1
Epha5
Nrxn1
Gene Expression z-Score
H
PIN Legend
Individual DEG Sets:
[1] Cytokines (n=10)
[2] Myelination (n=6)
[3] Oxidative Stress (n=7)
[4] Tissue Integrity (n=5)
Intersections:
Sets [1;3]: Cytokines;Oxidative Stress (n=3)
Sets [2;4]: Myelination;Tissue Integrity (n=1)
FLT4
PTGS2
BCL2
FAS
NTF3
FLT1
HGF
VEGFA
TGFB1
TNFRSF1A
CXCR4
ICAM1
JUN
IL1R1
CXCL12
CD4
AQP4
CD9
FN1
ITGA7
IL4RA
MOG
MAG
PLLP

**Supplemental Fig. 8. Hv1 deficiency reshapes the cortical transcriptional response to chronic TBI, converging on oxidative-stress and connected pathways. (A)** Unsupervised hierarchical clustering of group-level expression separates the four groups. **(B)** PLS-DA score plot showing genotype- and injury-resolved clustering of the four groups. **(C)** Per-pathway ssGSEA scores by group, with pathways ordered by hierarchical clustering into four modules; Hv1KO/TBI cortex shows the most divergent profile. **(D)** Volcano plot of differential expression for Hv1KO/TBI versus WT/TBI ($p < 0.05$): 103 DEGs (56 up, 47 down) of 674 genes. **(E)** Heatmap of the 103 DEGs across individual WT/TBI and Hv1KO/TBI samples (left), with the alluvial diagram (right) assigning most DEGs to the top five pathways (Oxidative Stress, Transcription and Splicing, Unfolded Protein Response, Carbohydrate Metabolism, Autophagy). **(F)** Panel-based GSEA on the Hv1KO/TBI-versus-WT/TBI ranking: positive enrichment (activation) of Oxidative Stress and Cytokines, and negative enrichment (suppression) of Myelination and Tissue Integrity. **(G)** Per-sample expression of representative leading-edge genes from the four GSEA pathways in (F). **(H)** Protein-protein interaction network (PIN) of genes from the four enriched pathways, and red dashed circles mark cross-pathway hub genes (BCL2, VEGFA, CXCL12, CD9, AQP4). Sample sizes: n=6 mice/group.

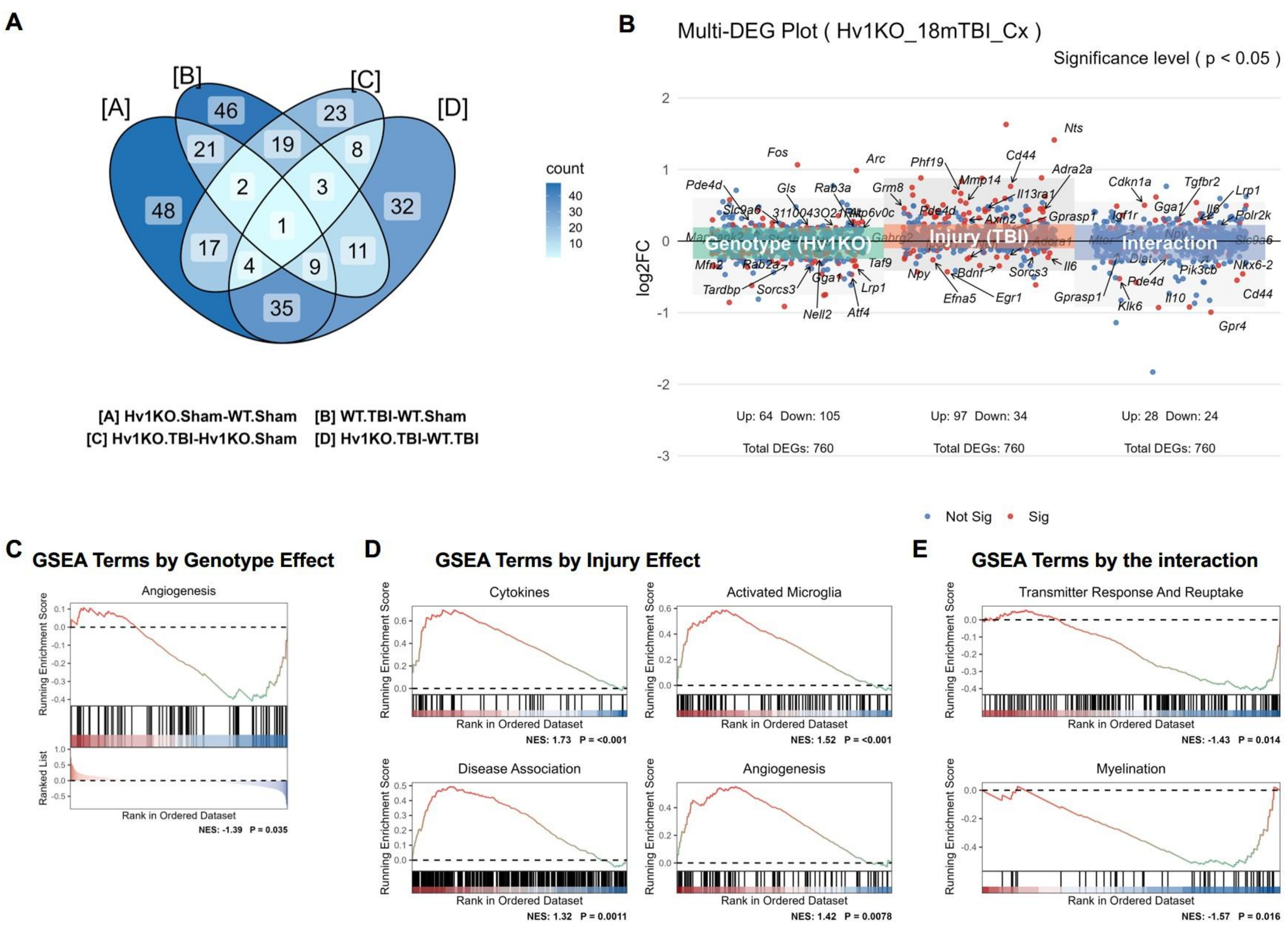


**Supplemental Fig. 9. Factor decomposition of the cortical Neuropathology panel (genotype × injury). (A)** Venn diagram of DEGs shared among four pairwise comparisons: [A] Hv1KO/Sham vs WT/Sham, [B] WT/TBI vs WT/Sham, [C] Hv1KO/TBI vs Hv1KO/Sham, [D] Hv1KO/TBI vs WT/TBI; shading indicates DEG count per region. **(B)** Multi-DEG plot of per-gene $\log_2$ fold change decomposed into genotype (Hv1KO vs WT), injury (TBI vs Sham), and interaction terms; red, significant ($p < 0.05$), blue, non-significant. Significant DEGs: genotype 169 (64 up, 105 down), injury 131 (97 up, 34 down), interaction 52 (28 up, 24 down). **(C)** GSEA on the genotype-effect ranking: negative enrichment (suppression) of "Angiogenesis". **(D)** GSEA on the injury-effect ranking: positive enrichment (activation) of "Cytokines", "Activated Microglia", "Disease Association", and "Angiogenesis". **(E)** GSEA on the interaction-effect ranking: negative enrichment (suppression) of "Transmitter Response and Reuptake" and "Myelination". Sample sizes: n=6 mice/group.

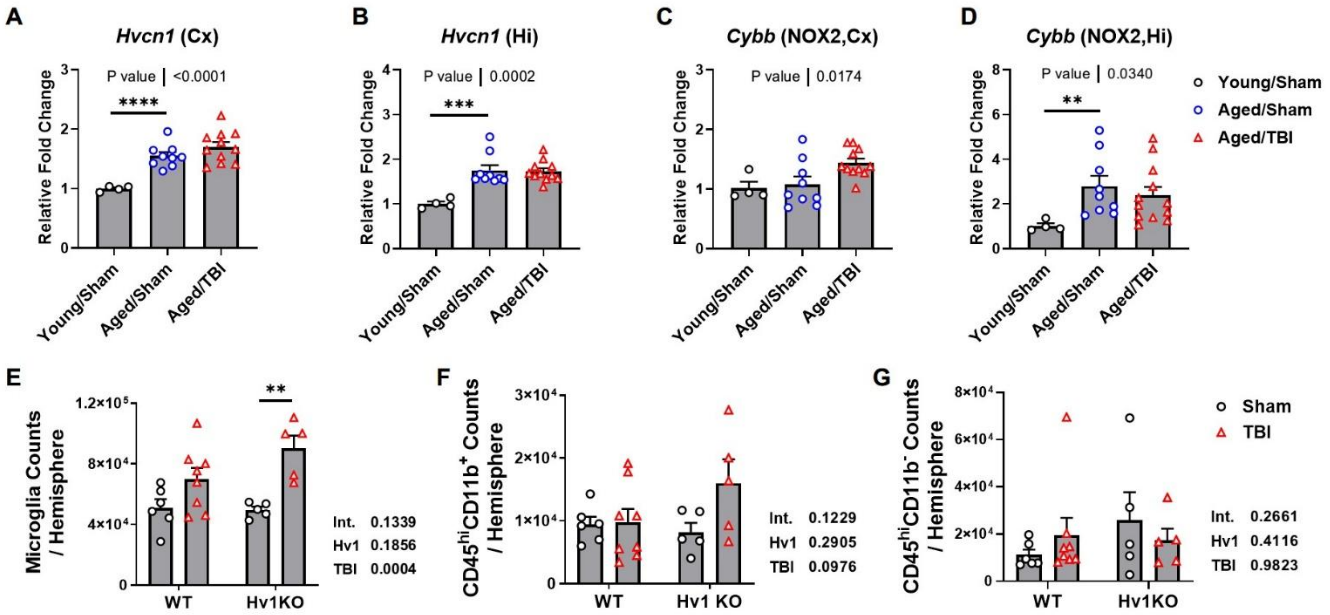


**Supplemental Fig. 10. Hv1 expression is age-dependent in the cortex and hippocampus, and its genetic ablation alters immune profiles in the brain after chronic TBI. (A-B)** The expression of Hvcn1 was assessed with age and TBI at mRNA level by qPCR in the cortex (A) and hippocampus (B) in wild-type C57BL6 mice of Young/Sham, Aged/Sham, and Aged/TBI. **(C-D)** The expression of Cybb (NOX2) was assessed with age and TBI at mRNA level by qPCR in the cortex (C) and hippocampus (D) in wild-type C57BL6 mice of Young/Sham, Aged/Sham, and Aged/TBI. **(E)** Increased microglia cells were detected in the brain of TBI mice by flow-cytometry compared to Sham groups. **(F-G)** Flow-cytometry data indicated no significant changes in infiltrated myeloid (F) and lymphocytes (G) across the four groups. Sample sizes for (A-D): Young/Sham n=4; Aged/Sham n=9; Aged/TBI n=11. Sample sizes for (E-G): WT/Sham n=6; Hv1KO/Sham n=5; WT/TBI n=8; Hv1KO/TBI n=5. Statistical analyses: Brown-Forsythe and Welch ANOVA tests with post-hoc Dunnett's T3 test were performed for (A-D); ); two-way ANOVA with Šídák-corrected simple-effects comparisons (E-G) with p-values representing interaction (Int.), *Hvcn1* knock-out (Hv1), and TBI main-effect shown beside each panel. *p<0.05, **p<0.01, ***p<0.001, ****p<0.0001.
