## Supplemental Tables for "Hv1 proton channel is essential for splenic antibacterial defense and immune homeostasis during the chronic phase of traumatic brain injury in male mice"

**Supplementary Information**

Supplemental Information includes six Supplemental tables and table legends.

**Supplemental Table 1. RT-qPCR primers information.**

| Gene | Assay ID | Target |
|---|---|---|
| *Escherichia coli* | Ba04646242_s1 | Escherichia coli |
| *Staphylococcus aureus* | Ba04646259_s1 | Staphylococcus aureus |
| *Total Bacteria 16s* | Ba04230899_s1 | 16S pan-bacterial Control |
| *Acot3* | Mm00652967_m1 | acyl-CoA thioesterase 3 |
| *Apoe* | Mm01307192_m1 | apolipoprotein E |
| *Batf3* | Mm01318274_m1 | basic leucine zipper transcription factor, ATF-like 3 |
| *Ccl5* | Mm01302427_m1 | chemokine (C-C motif) ligand 5 |
| *Cldn2* | Mm00516703_s1 | claudin 2 |
| *Clec7a* | Mm01183350_m1 | C-type lectin domain family 7, member a |
| *Ctss* | Mm01255859_m1 | cathepsin S |
| *Cybb* (NOX2) | Mm01287743_m1 | cytochrome b-245, beta polypeptide |
| *Cyp3a13* | Mm00484110_m1 | cytochrome P450, family 3, subfamily a, polypeptide 13 |
| *Duspy6* | Mm00518185_m1 | dual specificity phosphatase 6 |
| *Fcgr4* | Mm07300046_g1 | Fc receptor, IgG, low affinity IV |
| *Hgf* | Mm01135184_m1 | hepatocyte growth factor |
| *Hmox1* | Mm00516005_m1 | heme oxygenase 1 |
| *Hpgd* | Mm00515121_m1 | hydroxyprostaglandin dehydrogenase 15 (NAD) |
| *Hvcn1* | Mm01199507_m1 | hydrogen voltage-gated channel 1 |
| *Id2* | Mm00711781_m1 | inhibitor of DNA binding 2 |
| *Ifnb1* | Mm00439552_s1 | interferon beta 1, fibroblast |
| *Il17ra* | Mm00434214_m1 | interleukin 17 receptor A |
| *Il1b* | Mm00434228_m1 | interleukin 1 beta |
| *Il22* | Mm01226722_g1 | interleukin 22 |
| *Irf3* | Mm00516784_m1 | interferon regulatory factor 3 |
| *Irgm2* | Mm00546343_s1 | immunity-related GTPase family M member 2 |
| *Itga4* | Mm01277951_m1 | integrin alpha 4 |
| *Jak3* | Mm00439973_m1 | Janus kinase 3 |

| | | |
|---|---|---|
| *Lcn2* | Mm01324470_m1 | lipocalin 2 |
| *Mb21d1* (cGAS) | Mm01147496_m1 | Mab-21 domain containing 1 |
| *Muc2* | Mm01276676_m1 | mucin 2 |
| *Ocln* | Mm00500910_m1 | occludin |
| *Prkca* | Mm00440858_m1 | protein kinase C, alpha |
| *Reg3g* | Mm00441127_m1 | regenerating islet-derived 3 gamma |
| *Syk* | Mm01333032_m1 | spleen tyrosine kinase |
| *Tff3* | Mm00495590_m1 | trefoil factor 3, intestinal |
| *Tgm2* | Mm00436979_m1 | transglutaminase 2, C polypeptide |
| *Tjp1* | Mm01320638_m1 | tight junction protein 1 |
| *Tlr13* | Mm01233819_m1 | toll-like receptor 13 |
| *Tlr8* | Mm04209873_m1 | toll-like receptor 8 |
| *Tmem173* (STING) | Mm01158117_m1 | transmembrane protein 173 |
| *Tnf* | Mm00443258_m1 | tumor necrosis factor |
| *Tnfrsf1b* | Mm00441889_m1 | tumor necrosis factor receptor superfamily, member 1b |
| *GAPDH* | Mm99999915_g1 | glyceraldehyde-3-phosphate dehydrogenase |

All the primers were purchased from ThermoFisher/Invitrogen.

**Supplemental Table 2. ssGSEA scores of Myeloid Innate Immunity panel are significantly altered by Hv1 KO, exhibiting distinct pathway-wise responses to TBI in mice spleen.**

| Pathway | Gene | Injury | Gene x Injury |
|---|---|---|---|
| Angiogenesis | 0.0387 | 0.0076 | 0.0070 |
| Antigen Presentation | 0.1700 | 0.0318 | 0.3135 |
| Cell Cycle and Apoptosis | 0.0294 | 0.7098 | 0.6767 |
| Cell Migration and Adhesion | 0.0552 | 0.0482 | 0.2295 |
| Chemokine signaling | 0.4742 | 0.1716 | 0.8691 |
| Complement Activation | 0.0041 | 0.1985 | 0.3935 |
| Cytokine Signaling | 0.0001 | 0.1024 | 0.0262 |
| Differentiation and Maintenance of Myeloid Cells | 0.0002 | 0.5867 | 0.4461 |
| ECM remodeling | 0.0247 | 0.8595 | 0.7611 |
| Fc Receptor Signaling | 0.0025 | 0.0306 | 0.0144 |
| Growth Factor Signaling | 0.0000 | 0.0307 | 0.0042 |
| Interferon Signaling | 0.0059 | 0.0612 | 0.5606 |
| Lymphocyte activation | 0.0000 | 0.0271 | 0.0632 |
| Metabolism | 0.9285 | 0.6384 | 0.1824 |
| Pathogen Response | 0.0001 | 0.0768 | 0.0062 |
| T-cell Activation and Checkpoint Signaling | 0.0110 | 0.3114 | 0.1765 |
| TH1 Activation | 0.0017 | 0.8763 | 0.3355 |
| TH2 Activation | 0.0881 | 0.3274 | 0.3689 |
| TLR signaling | 0.0004 | 0.1323 | 0.5147 |

P values corresponding to the factor of Hv1 KO, TBI, and their interaction based on the two-way ANOVA analysis for each pathway. WT/Sham, n=6; Hv1KO/Sham, n=4; WT/TBI, n=6; Hv1KO/TBI, n=4.

**Supplemental Table 3. ssGSEA scores of Myeloid Innate Immunity panel are predominantly altered by Hv1 KO with TBI in mice lung.**

| Pathway | Gene | Injury | Gene x Injury |
|---|---|---|---|
| Angiogenesis | 0.0457 | 0.5021 | 0.9554 |
| Antigen Presentation | 0.1198 | 0.9481 | 0.1863 |
| Cell Cycle and Apoptosis | 0.0001 | 0.4574 | 0.2211 |
| Cell Migration and Adhesion | 0.1899 | 0.1710 | 0.1801 |
| Chemokine signaling | 0.0000 | 0.7001 | 0.7259 |
| Complement Activation | 0.0000 | 0.0425 | 0.1096 |
| Cytokine Signaling | 0.0751 | 0.2938 | 0.9782 |
| Differentiation and Maintenance of Myeloid Cells | 0.0018 | 0.7853 | 0.1029 |
| ECM remodeling | 0.1340 | 0.1430 | 0.9839 |
| Fc Receptor Signaling | 0.0190 | 0.3739 | 0.8547 |
| Growth Factor Signaling | 0.0012 | 0.6723 | 0.7545 |
| Interferon Signaling | 0.3606 | 0.8087 | 0.2401 |
| Lymphocyte activation | 0.6411 | 0.9050 | 0.3580 |
| Metabolism | 0.1752 | 0.5949 | 0.7943 |
| Pathogen Response | 0.0371 | 0.1912 | 0.2104 |
| T-cell Activation and Checkpoint Signaling | 0.0769 | 0.6519 | 0.5591 |
| TH1 Activation | 0.0001 | 0.1913 | 0.8199 |
| TH2 Activation | 0.6265 | 0.1252 | 0.3751 |
| TLR signaling | 0.0004 | 0.4311 | 0.3529 |

P values corresponding to the factor of Hv1 KO, TBI, and their interaction based on the two-way ANOVA analysis of ssGSEA scores per pathway. WT/Sham, n=4; Hv1KO/Sham, n=6; WT/TBI, n=5; Hv1KO/TBI, n=6.

**Supplemental Table 4. ssGSEA scores of Myeloid Innate Immunity panel are predominantly altered by Hv1 KO with TBI in mice liver.**

| Pathway | Gene | Injury | Gene x Injury |
|---|---|---|---|
| Angiogenesis | 0.2266 | 0.4594 | 0.0083 |
| Antigen Presentation | 0.1411 | 0.1469 | 0.0229 |
| Cell Cycle and Apoptosis | 0.1288 | 0.4024 | 0.8051 |
| Cell Migration and Adhesion | 0.3140 | 0.7231 | 0.1201 |
| Chemokine signaling | 0.4755 | 0.4974 | 0.0165 |
| Complement Activation | 0.0052 | 0.7073 | 0.0185 |
| Cytokine Signaling | 0.2362 | 0.1856 | 0.0229 |
| Differentiation and Maintenance of Myeloid Cells | 0.1839 | 0.4107 | 0.9067 |
| ECM remodeling | 0.5625 | 0.7665 | 0.1978 |
| Fc Receptor Signaling | 0.0698 | 0.1416 | 0.0157 |
| Growth Factor Signaling | 0.1310 | 0.5000 | 0.0102 |
| Interferon Signaling | 0.8048 | 0.6366 | 0.0175 |
| Lymphocyte activation | 0.0005 | 0.4701 | 0.0059 |
| Metabolism | 0.0187 | 0.5419 | 0.0044 |
| Pathogen Response | 0.0002 | 0.6101 | 0.0363 |
| T-cell Activation and Checkpoint Signaling | 0.0132 | 0.2915 | 0.4332 |
| TH1 Activation | 0.6847 | 0.7269 | 0.2083 |
| TH2 Activation | 0.2679 | 0.4662 | 0.5018 |
| TLR signaling | 0.6350 | 0.0949 | 0.2335 |

P values corresponding to the factor of Hv1 KO, TBI, and their interaction based on the two-way ANOVA analysis of ssGSEA scores per pathway. WT/Sham, n = 6; Hv1KO/Sham, n = 6; WT/TBI, n = 6; Hv1KO/TBI, n = 6.

**Supplemental Table 5. Cell-Type enrichment statistics in mouse spleen (A), lung (B), and liver (C).**

| **A** | **Spleen** | | |
|---|---|---|---|
| **Cell Type** | **Gene** | **Injury** | **Gene x Injury** |
| Neutrophils | 0.0000 | 0.0218 | 0.0022 |
| CD45 | 0.0710 | 0.0163 | 0.0279 |
| Macrophages | 0.0096 | 0.6802 | 0.0650 |
| DC | 0.2685 | 0.6446 | 0.1126 |
| Cytotoxic cells | 0.3252 | 0.2633 | 0.2815 |
| B-cells | 0.0000 | 0.9006 | 0.3449 |
| Mast cells | 0.0000 | 0.4109 | 0.3635 |
| NK CD56dim cells | 0.1525 | 0.1781 | 0.6258 |
| Exhausted CD8 | 0.0009 | 0.7699 | 0.8064 |

| **B** | **Lung** | | |
|---|---|---|---|
| **Cell Type** | **Gene** | **Injury** | **Gene x Injury** |
| Cytotoxic cells | 0.0021 | 0.0123 | 0.0323 |
| DC | 0.0034 | 0.1955 | 0.1752 |
| Neutrophils | 0.0008 | 0.8816 | 0.2503 |
| Exhausted CD8 | 0.0437 | 0.2935 | 0.3559 |
| Macrophages | 0.0002 | 0.6326 | 0.3938 |
| NK CD56dim cells | 0.1932 | 0.0920 | 0.4300 |
| Mast cells | 0.1260 | 0.6954 | 0.6531 |
| CD45 | 0.0161 | 0.5656 | 0.9196 |
| B-cells | 0.0479 | 0.2192 | 0.9435 |

**C** **Liver**

| Cell Type | Gene | Injury | Gene x Injury |
|---|---|---|---|
| CD45 | 0.2452 | 0.8717 | 0.0207 |
| Macrophages | 0.1023 | 0.2787 | 0.0380 |
| Neutrophils | 0.0023 | 0.2612 | 0.0397 |
| DC | 0.1379 | 0.1806 | 0.1110 |
| Cytotoxic cells | 0.9859 | 0.4444 | 0.1653 |
| B-cells | 0.0699 | 0.5232 | 0.1830 |

P values corresponding to the factor of Hv1 KO, TBI, and their interaction based on the two-way ANOVA analysis for each pathway. Spleen: WT/Sham N=6; Hv1KO/Sham N=4; WT/TBI N=6; Hv1KO/TBI N=4. Lung: WT/Sham N=4; Hv1KO/Sham N=6; WT/TBI N=5; Hv1KO/TBI N=6. Liver: WT/Sham N=6; Hv1KO/Sham N=6; WT/TBI N=6; Hv1KO/TBI N=6.

**Supplemental Table 6. ssGSEA scores of the Neuropathology panel are predominantly altered by Hv1 KO and chronic TBI in the mouse cortex.**

| Pathway | Gene | Injury | Gene x Injury |
|---|---|---|---|
| Activated Microglia | 0.5991 | 0.7273 | 0.6956 |
| Angiogenesis | 0.0074 | 0.0057 | 0.0029 |
| Apoptosis | 0.1790 | 0.0339 | 0.8985 |
| Autophagy | 0.7370 | 0.1231 | 0.1030 |
| Axon and Dendrite Structure | 0.0646 | 0.9658 | 0.8705 |
| Carbohydrate Metabolism | 0.3469 | 0.1567 | 0.8395 |
| Chromatin Modification | 0.3873 | 0.5242 | 0.9143 |
| Cytokines | 0.0229 | 0.0239 | 0.0326 |
| Growth Factor Signaling | 0.7813 | 0.0130 | 0.1696 |
| Lipid Metabolism | 0.0091 | 0.0058 | 0.2248 |
| Matrix Remodeling | 0.2590 | 0.0036 | 0.6086 |
| Myelination | 0.3884 | 0.1385 | 0.4411 |
| Neural Connectivity | 0.2788 | 0.7544 | 0.6493 |
| Neuronal Cytoskeleton | 0.3519 | 0.0222 | 0.7811 |
| Oxidative Stress | 0.0123 | 0.0199 | 0.6575 |
| Tissue Integrity | 0.0224 | 0.0192 | 0.0093 |
| Transcription and Splicing | 0.2950 | 0.3433 | 0.3926 |
| Transmitter Release | 0.3516 | 0.9088 | 0.9839 |
| Transmitter Response and Reuptake | 0.2627 | 0.3028 | 0.9489 |

P values corresponding to the factor of Hv1 KO, TBI, and their interaction based on the two-way ANOVA analysis of ssGSEA scores per pathway. WT/Sham, n=6; Hv1KO/Sham, n=6; WT/TBI, n=6; Hv1KO/TBI, n=6.
